# Investigating the role of lipid genes in liver disease using fatty liver models of alcohol and high fat in zebrafish (*Danio rerio*)

**DOI:** 10.1101/2023.04.14.536511

**Authors:** Fathima Shihana, Pradeep Manuneedhi Cholan, Stuart Fraser, Stefan H Oehlers, Devanshi Seth

## Abstract

**Background:** Accumulation of lipid in the liver is the first hallmark of both alcohol-related liver disease (ALD) and non-alcohol-related fatty liver disease (NAFLD). Recent studies indicate that specific mutations in lipid genes confer risk and might influence disease progression to irreversible liver cirrhosis. This study aimed to understand the function/s of lipid risk genes driving disease development in zebrafish genetic models of alcohol- and non-alcohol related fatty liver.

**Methods:** We used zebrafish larvae to investigate the effect of alcohol and high fat to model fatty liver and tested the utility of this model to study lipid risk gene functions. CRISPR/Cas9 gene editing was used to create knockdowns in 5 days post-fertilization zebrafish larvae for the available orthologs of human cirrhosis risk genes (*pnpla3, faf2, tm6sf2*). To establish fatty liver models, larvae were exposed to ethanol and a high fat diet (HFD) consisting of chicken egg yolk. Changes in morphology (imaging), survival, liver injury (biochemical tests, histopathology), gene expression (qPCR) and lipid accumulation (dye specific live imaging) were analysed across treatment groups to test the functions of these genes.

**Results:** Exposure of 5-day post-fertilization (dpf) WT larvae to 2% ethanol or HFD for 48 hours developed measurable hepatic steatosis. CRISPR-Cas9 genome editing depleted *pnpla3, faf2* and *tm6sf2* gene expression in these CRISPR knockdown larvae (crispants). Knockdown significantly increased effects of ethanol and HFD toxicity by increasing hepatic steatosis and hepatic neutrophil recruitment ≥2-fold in all three crispants. Furthermore, ethanol or HFD exposure significantly altered the expression of genes associated with ethanol metabolism (*cyp2y3*) and lipid metabolism-related gene expression, including *atgl* (triglyceride hydrolysis), *axox1, echs1* (fatty acid β-oxidation), *fabp10a* (transport), *hmgcra* (metabolism), *notch1* (signaling) and *srebp1* (lipid synthesis), in all three *pnpla3, faf2* and *tm6sf2* crispants. Nile Red staining in all three crispants revealed significantly increased lipid droplet size and triglyceride accumulation in the livers following exposure to ethanol or HFD.

**Conclusions:** We identified roles for *pnpla3, faf2* and *tm6sf2* genes in triglyceride accumulation and fatty acid oxidation pathways in a zebrafish larvae model of fatty liver.

## Introduction

Fatty liver diseases driven by lifestyle factors, such as risky drinking and dietary excess, are predicted to escalate in future adding to the already mounting global liver disease burden. Accumulation of lipids (fat) in the liver is the first hallmark of both alcohol-related liver disease (ALD) and non-alcohol-related fatty liver disease (NAFLD). This persistent excess of lipid causes lipotoxicity, oxidative stress, and inflammation leading to fibrosis, cirrhosis and end-stage liver disease.

Fatty liver, the first step in chronic liver diseases, ALD and NAFLD, is characterised by lipid accumulation in more than 90% of chronic heavy drinking.[1] However, what begins as accumulation of fat in the liver progresses to inflammation, fibrosis and ultimately irreversible cirrhosis in up to 30% of excessive drinkers and NAFLD patients.[2] Abnormal lipid accumulation in hepatic lipid droplets (hepatic steatosis) is a recognised cause of prevalent metabolic diseases such as metabolic syndrome or obesity.[3] Lipid droplets are increasingly recognized as a defining characteristic of ALD [4] and NAFLD and also as having important non-pathological roles in cell signaling. Lipid droplets are highly regulated by the hundreds of proteins that coat the lipid droplet surface and control lipid trafficking and flux, hence investigating the risk genes associated with lipid droplets/biology will allow better understanding of their roles in disease development.[5, 6]

Recent studies indicate that specific gene mutations confer risk and might influence disease progression in some people to irreversible liver cirrhosis.[7-10] Our recent Genome-Wide Association Study (GWAS) in the GenomALC cohort of thousands of drinkers reported novel (*FAF2, SERPINA1*) and previously reported (*PNPLA3, SUGP1/TM6SF2, HSD17B13*) associations of single nucleotide polymorphisms (SNPs) with risk of alcohol- and NAFLD-related cirrhosis.[7, 11] Intriguingly, the novel *FAF2* variant, along with *PNPLA3, HSD17B13* and *SUGP1/TM6SF2* SNPs, are all implicated in lipid biology.[12-14] Recognition of lipids playing a central role in liver disease pathology is increasing[15], and lipid metabolism genes and related molecular pathways are potential therapeutic targets.

Zebrafish have proven to be an important tool for the high-throughput screening of drugs for hepatotoxicity[16-19] and as models of liver injury, including alcohol and non-alcohol-related liver diseases.[20-23] Furthermore, the fundamental physiologic processes, genetic mutations and manifestations of pathogenic responses to environmental insults exhibit many similarities with humans,[19, 24] and the larval transparency of the zebrafish allows real-time imaging of hepatic biology.[24, 25] Thus this study deployed zebrafish models of alcohol- and non-alcohol-related fatty liver with an aim to understand the function/s of lipid risk genes driving disease development.

## Materials and Methods

### Zebrafish strains and housing

Adult zebrafish were housed at the Centenary Institute under Sydney Local Health District AEC Approval 17–036. Wild-type (WT) zebrafish larvae were produced through the natural spawning of adult males and females from an AB genetic background. The Tg(lyzC:gfp)^nz117^ strain was used to visualise neutrophils. Zebrafish embryos were incubated in dark at 28°C. The E3 media with methylene blue was changed daily to reduce microbial growth.

### gRNA synthesis, CRISPR injections and creation of zebrafish larvae ‘crispants’

Guide RNA (gRNA) was synthesized for *pnpla3, faf2* and *tm6sf2* gene targets (see Table 1 for primers [26] and probes) as described in the manual of the manufacturer (Precision gRNA Synthesis Kit, Thermo Fisher Scientific). A 1:1 solution of gRNA and 500 µg/mL of Cas9 nuclease V3 (Integrated DNA Technology Iowa 52241, USA) was prepared with phenol red dye (Sigma, P0290). Freshly laid zebrafish eggs were collected from breeding tanks and 1 nL solution mix was injected using a FemtoJet 4i (Eppendorf) into the yolk before the first cellular division.

**Table 1:**
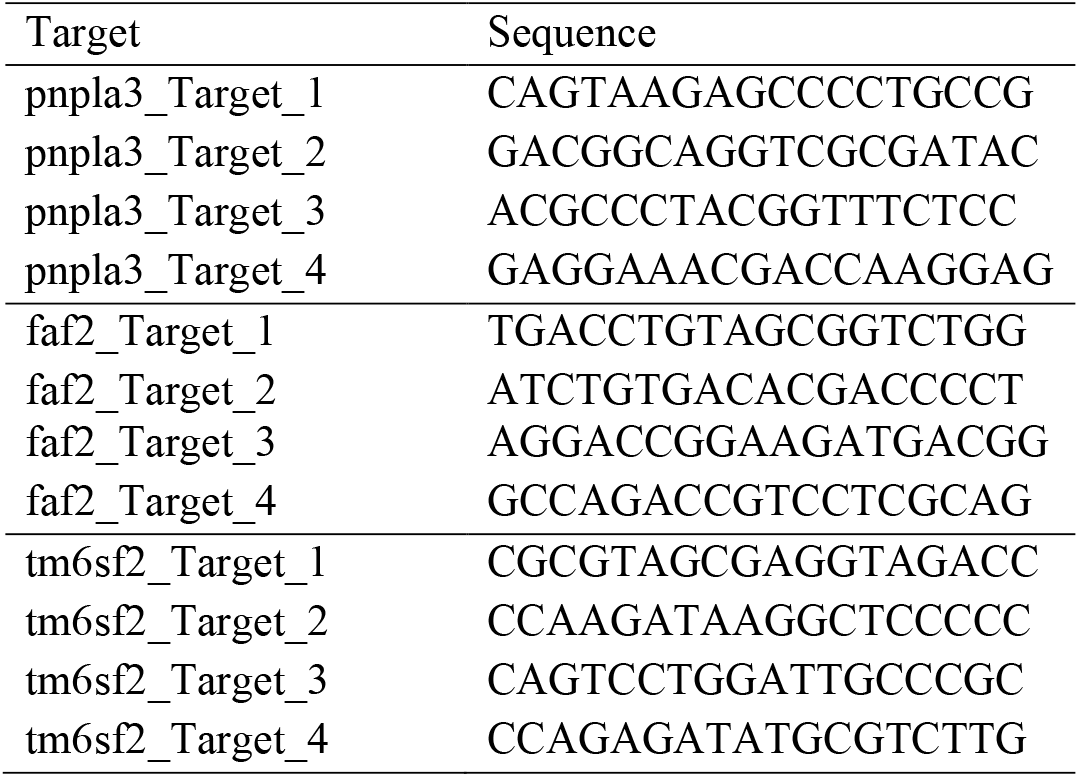
Zebrafish primer sequences for gRNA

### Models of EtOH and HFD diet-induced fatty liver

At 5 days post fertilisation (dpf) 40-50 larvae were exposed to ethanol (EtOH, concentrations as stated usually 2% v/v) for 48 hours in Petri dishes containing E3/methylene blue media.

A high fat diet (HFD) solution was prepared using boiled chicken egg yolk (0.5 ml) in 50 ml E3 solution. The solution was diluted to make a final concentration of 0.05% of emulsified chicken egg yolk.[27] Larvae (5 dpf, n=50-60) were exposed to E3 solution containing HFD for 48 hours and the media was changed on day 6.

### Oil Red O staining

Whole mount Oil Red O staining was performed on larvae as previously described.[28] Briefly, whole larvae were fixed with 4% paraformaldehyde, washed with PBS, infiltrated with a graded series of propylene glycol, and stained with 0.5% Oil Red O in 100% propylene glycol overnight. Stained larvae were washed with decreasing concentrations of propylene glycol followed by several rinses with PBS and transferred to 3% methylcellulose for imaging. Oil Red O staining was quantified in Fiji (ImageJ) by adjusting the colour threshold to eliminate the non-red signal. The image was then converted to a binary mask, and the liver region was selected to measure the number and area of particles.[27]

### Neutrophil counts

Transgenic Tg(lyzC:gfp)^nz117^ larvae were anesthetized with tricaine and mounted in 3% methylcellulose. Larvae were imaged using a Leica M205FA fluorescent microscope for GFP-expressing neutrophils. Fiji (ImageJ) software was used to quantify neutrophil fluorescent pixel counts within the area of the liver.

### Gene expression using quantitative PCR (qPCR)

10–15 zebrafish larvae per treatment group were pooled and lysed using a 23-gauge needle in Trizol LS for RNA extraction. The primers for reverse transcription-polymerase chain reaction (RT-PCR) are listed in Table 2. cDNA was synthesized using a High-capacity reverse transcription kit (ThermoFisher Scientific, 4368814). qPCR was carried out using Power UP SYBR green master mix (ThermoFisher Scientific, 4385610) on a CFX96 Real-Time system (BioRad).

**Table 2:**
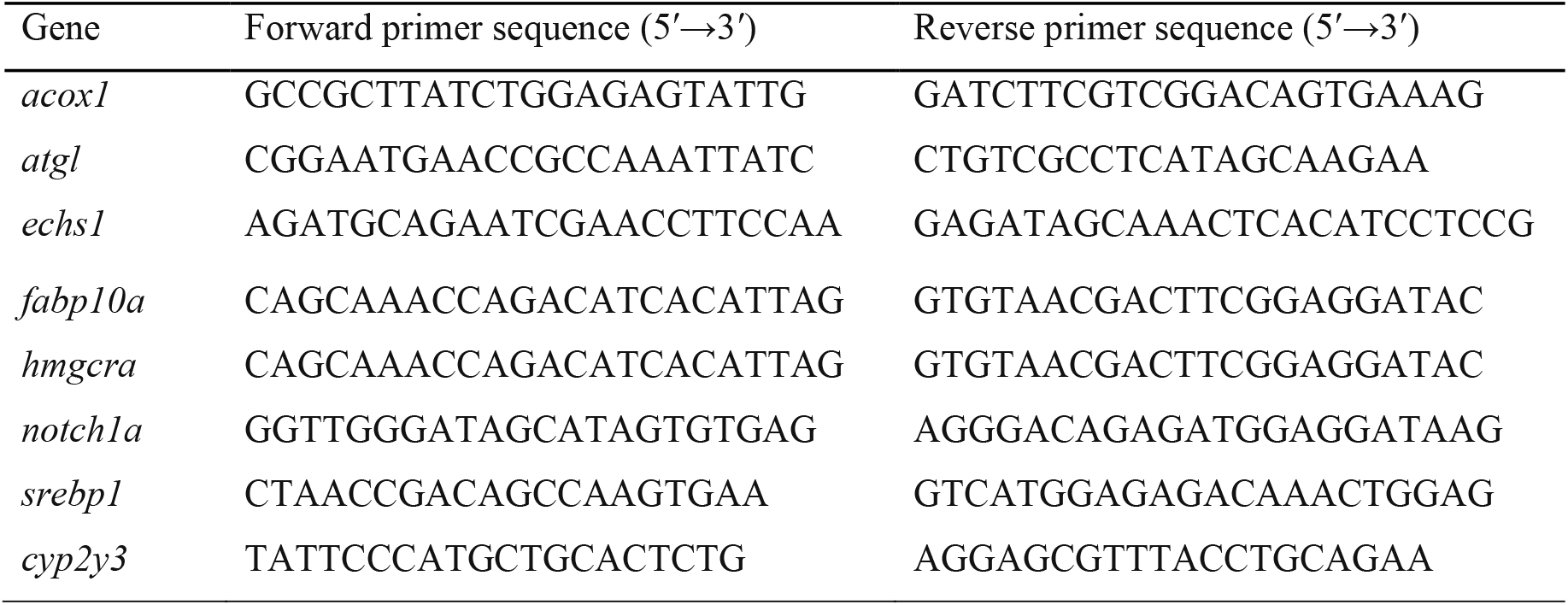
Zebrafish primer sequences for RT-PCR

### Nile Red and Nile Blue Staining and confocal microscopic imaging

A set of 5-8 zebrafish larvae were stained in E3 solution (300 *μ*L) containing Nile Red (300 nM) and Nile Blue (5 µM) for 30 minutes [29]. Zebrafish larvae were anesthetized using tricaine. For imaging, larvae were immobilised using 1% w/v low melting point (LMP) agarose dissolved in E3 solution. Two-three zebrafish larvae were embedded in LMP within a 35 mm glass bottom dish (Mattek) and covered with E3/tricaine.

High-resolution images were taken using a Leica SP8 confocal microscope coupled with the LasX software package. Nile Red and Nile Blue were excited using the 488 nm and 643 nm lasers respectively.

### Statistical analysis

Statistical testing was carried out using GraphPad Prism 9, with correction for multiple comparisons or Student’s *t*-tests as appropriate. All data are representative of at least three experimental each with 10-15 biological replicates. Outliers were excluded using ROUT with Q = 1% in Prism. Data are expressed as median with IQR (5,95). Every data point represents a single larva unless otherwise noted. One-way ordinary ANOVA with a Sidak multiple comparison test with single pooled variance was used to compare the selected group in the dataset.

## Results

### Exposure to EtOH increases fat but has no effect on neutrophilic inflammation in zebrafish larval livers

Treatment with 2% EtOH for 48 hours showed near 100% survival, while 25% and 100% of larvae died at 2.5% and 4% EtOH, respectively (Supplementary Figure 1A). We selected 2% EtOH for 48 hours as our optimal dose to conduct further experiments. Morphological changes, mainly upward curvature of the trunk and tail were seen at 2% EtOH dose (Supplementary Figure 1B). No other significant morphological abnormalities were noted with this treatment. ORO staining showed significantly increased hepatic lipids in the EtOH exposed larvae compared to controls (Supplementary Fig 1C). There was no significant difference in hepatic neutrophil count caused by this alcohol exposure challenge (Supplementary Fig 1D).

### Knockdown characteristics of pnpla3, faf2 and tm6sf2 crispants

Knockdown was confirmed by measuring *pnpla3, faf2*, and *tm6sf2* gene expression in CRISPR-Cas9 treated larvae. CRISPR targeting of *pnpla3, faf2*, and *tm6sf2* significantly decreased the expression of *pnpla3* (71%), *faf2* (89%), and *tm6sf2* (83%), respectively (Figure 1A). There was a marginally reduced survival by 7 dpf in *pnpla3* (10%), *faf2* (20%) and *tm6sf2* (4%) crispants but these differences were not statistically significant from scramble control cohorts (Figure 1B). Exposure to 2% EtOH, but not HFD, reduced survival in the crispants compared to their scramble controls, however, this was not significant from treatment control (Supplementary Figure 2A). When exposed to EtOH, the *pnpla3, faf2 and tm6sf2* crispants, compared to their respective scramble controls, more larvae showed overall morphological deficits such as body curvature (∼40%), edema (∼20%) and reduced yolk sac consumption (∼35%) (Figure 1C, Supplementary Figure 2B). No other morphological defect was observed in HFD diet exposed crispants (Supplementary Figure 2C & 2D).

**Figure 1:**
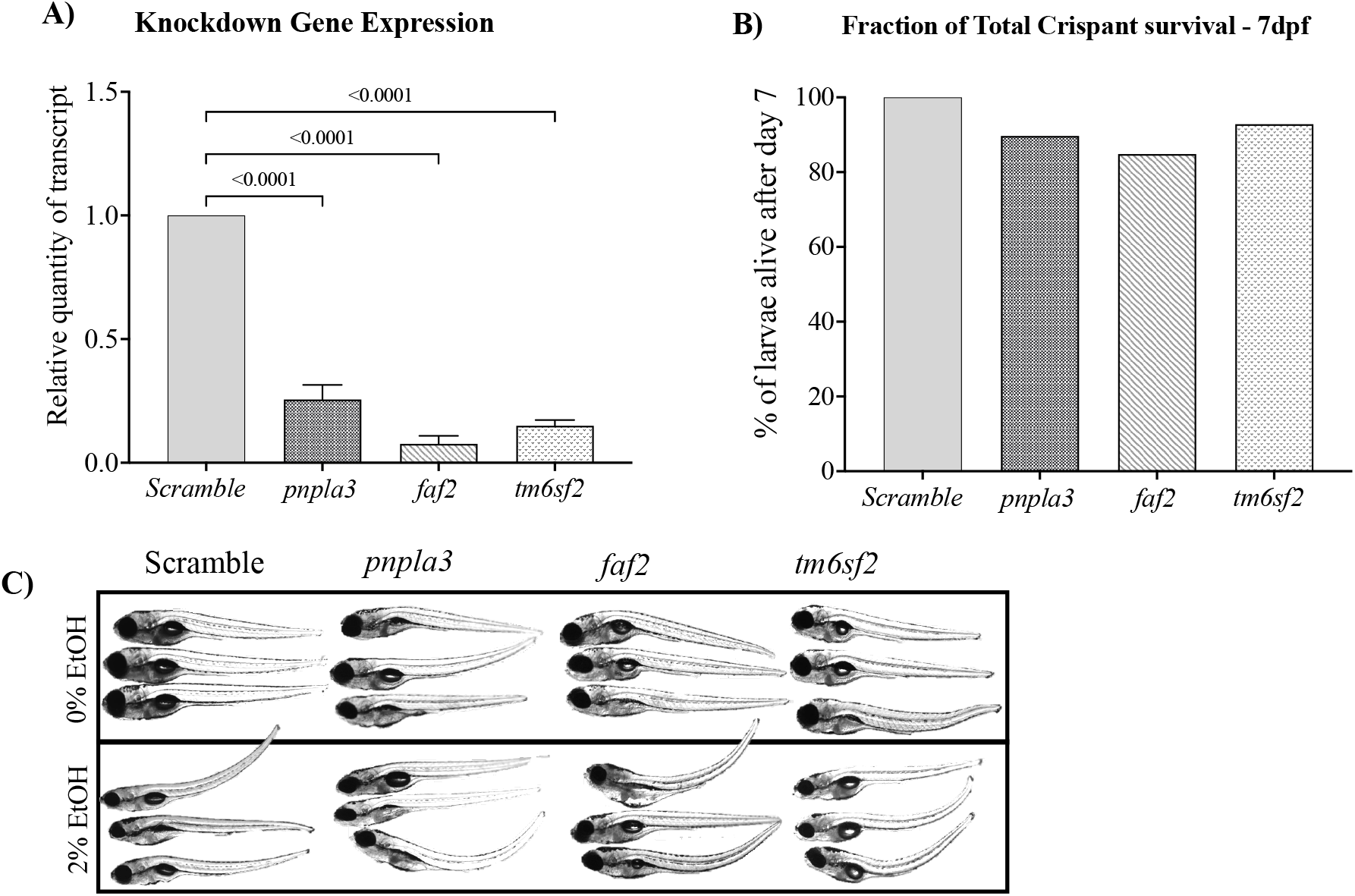
Survival and gene expression analysis in crispant zebrafish larvae. A) gene expression B) mortality *of pnpla3, faf2* and *tm6sf2*. Each bar is with three repeated experiments (clutch, 15-20 larvae in each clutch). * Indicates *p* < 0.05 crispant-fold change compared to scramble control as 1 using non-parametric Mann-Whitney test. C) Brightfield images from live Tg(lyzC:gfp)^nz117^ showing a range of developmental morphological abnormalities in *pnpla3, faf2 and tm6sf2* crispants with 2% EtOH exposure.

### Increased liver neutrophilic inflammation and fat in the crispants following EtOH or HFD exposure

Knockdown of *pnpla3, faf2* and *tm6sf2* did not increase hepatic neutrophil infiltration (Figure 2A). However, when exposed to 2% EtOH, hepatic neutrophil infiltration increased in *pnpla3* (3-fold), *faf2* (2-fold) and *tm6sf2* (3-fold) crispants compared to scramble control. Hepatic neutrophil infiltration also increased ∼3-fold across all crispants compared to scramble control exposed to HFD (Figure 2A).

**Figure 2:**
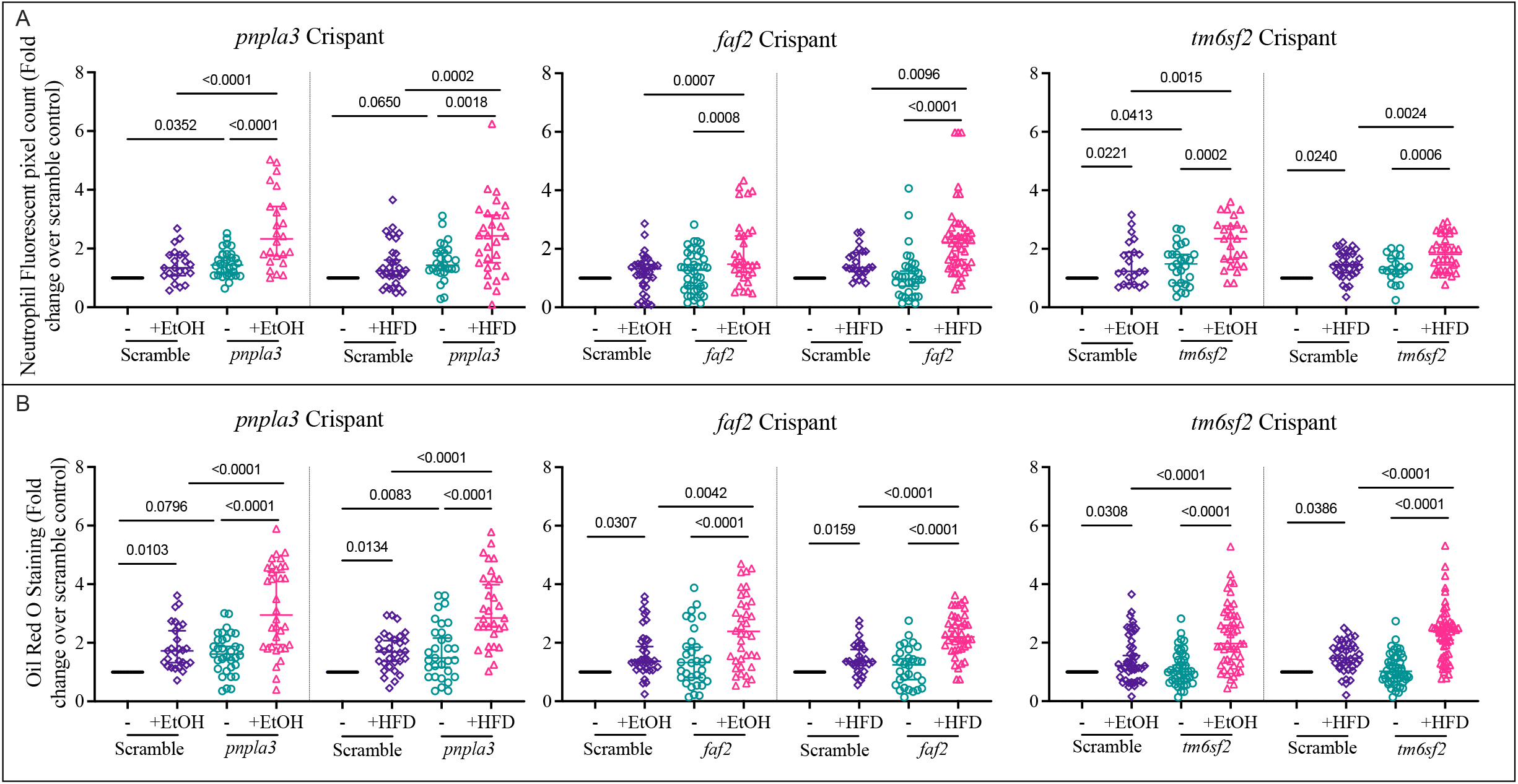
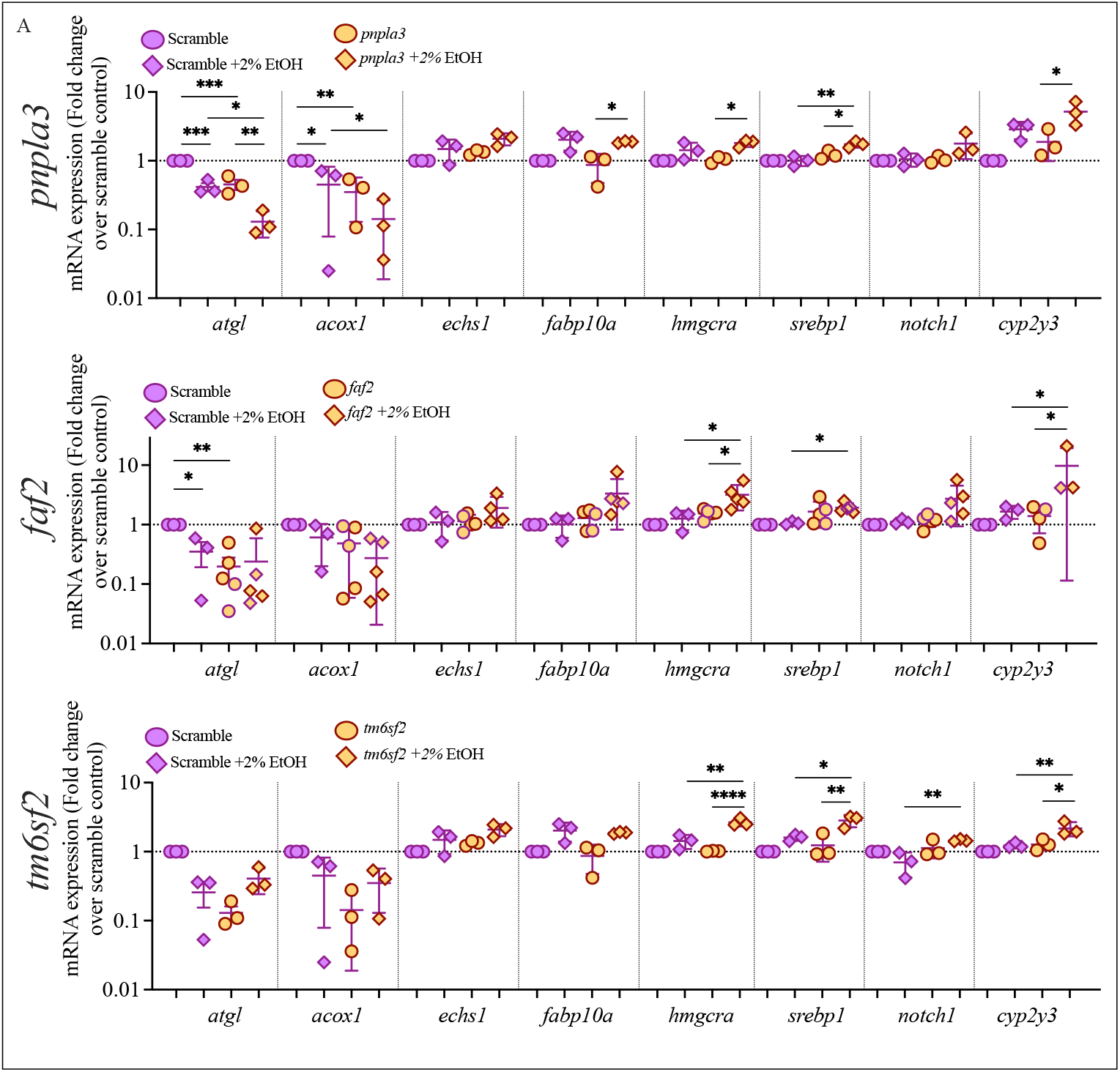
Ethanol or HFD exposure increased liver neutrophil and lipid accumulation. A) ORO staining showing lipid deposit in the zebrafish larva liver. B) Quantification of neutrophil recruitment in liver (A) quantification of lipid accumulation (B) using Oil Red O staining after exposure to 2% EtOH or HFD for 48 hours *in pnpla3, faf2* and *tm6sf2* crispants. Y- axis represents the foldchange over scramble control. Ordinary one-way ANOVA test was used to compare between respective groups.

Knockdown of *pnpla3*, but not *faf2* and *tm6sf2*, caused an increase in hepatic ORO lipid staining (Figure 2B). Hepatic ORO lipid staining was increased in *pnpla3, faf2* and *tm6sf2* crispants in both EtOH and HFD exposure compared to their respective exposed scramble controls (Figure 2B). Representative images showing increased ORO staining in the liver in all knockdowns compared to scramble and with treatments versus controls are shown in Supplementary Figure 3.

### Gene expression in crispants with EtOH or HFD exposure

*Pnpla3, faf2* and *tm6sf2* crispants showed significantly changed expression of several genes involved in lipid biology and alcohol metabolism (Figure 3; for more details see Supplementary Figure 3). *Acox1* (triglyceride (TG) hydrolysis) and *atgl* (fatty acid oxidation) genes were significantly downregulated in all crispants versus their respective scramble controls. Exposure to EtOH or HFD further downregulated the expression of these genes in some pairings of gene knockdown and treatment compared to either unexposed knockdown or exposed scrambled control (Figure 3), but these effects were not seen for the majority of conditions analysed.

**Figure 3:**
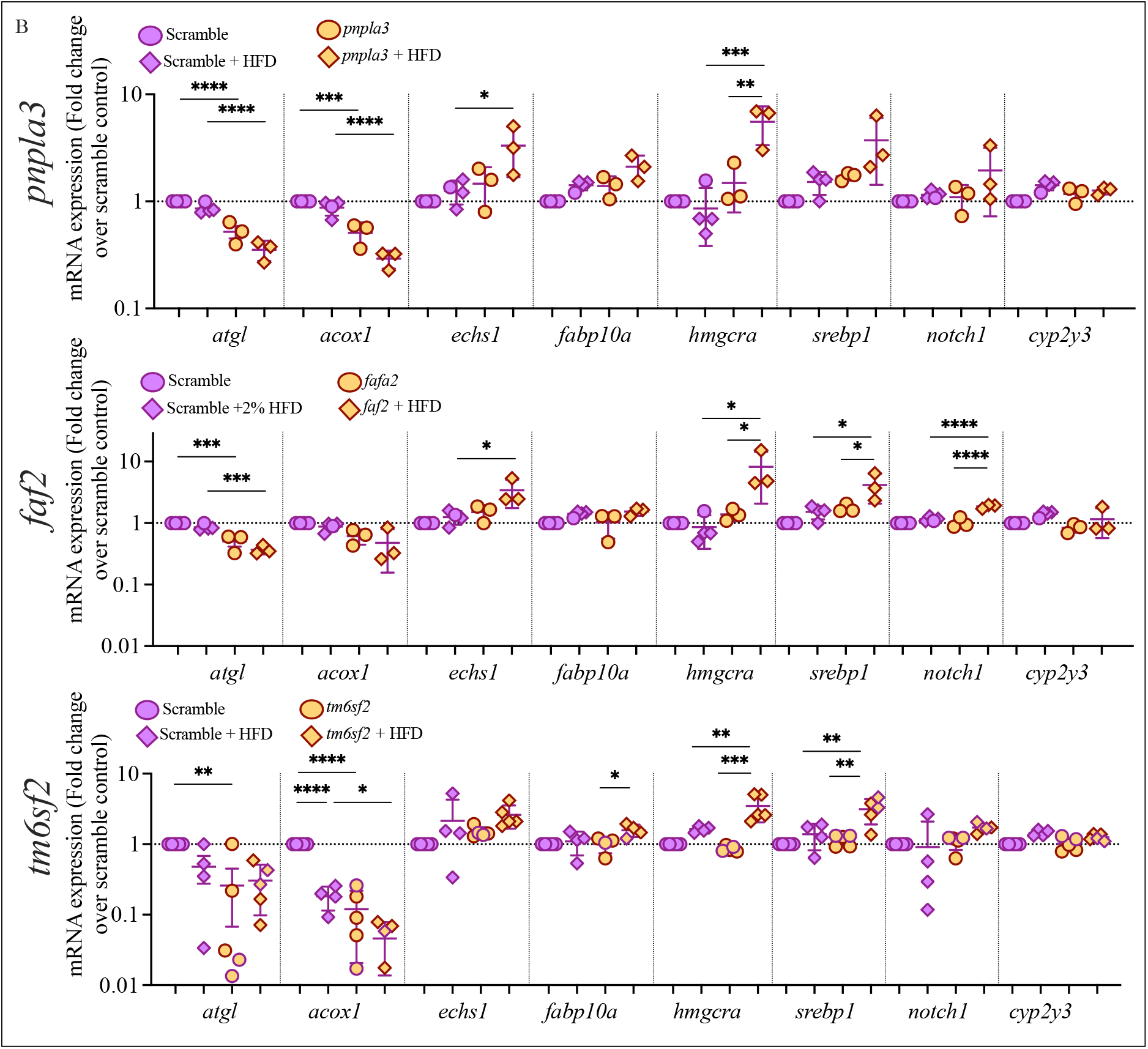
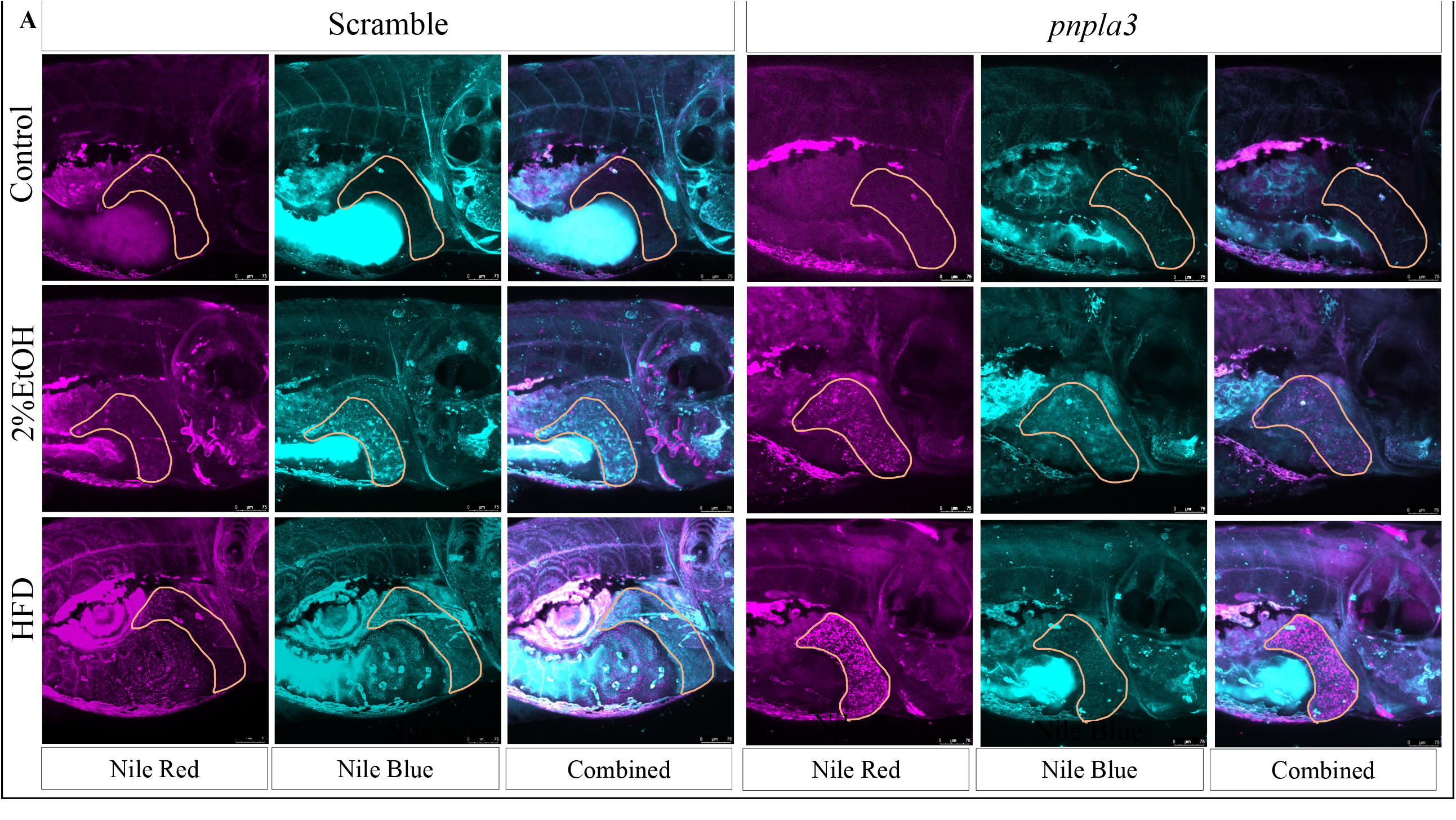
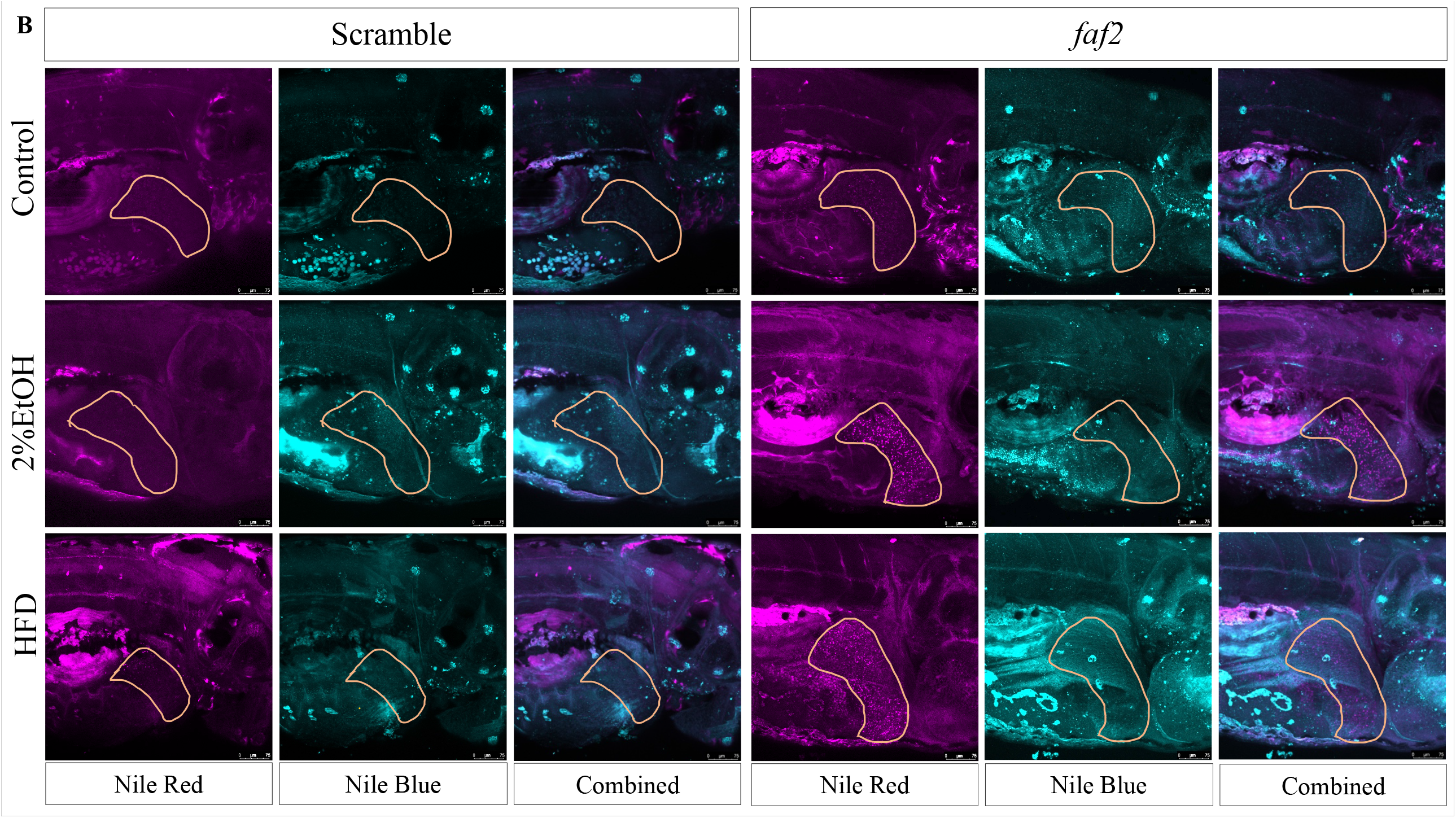
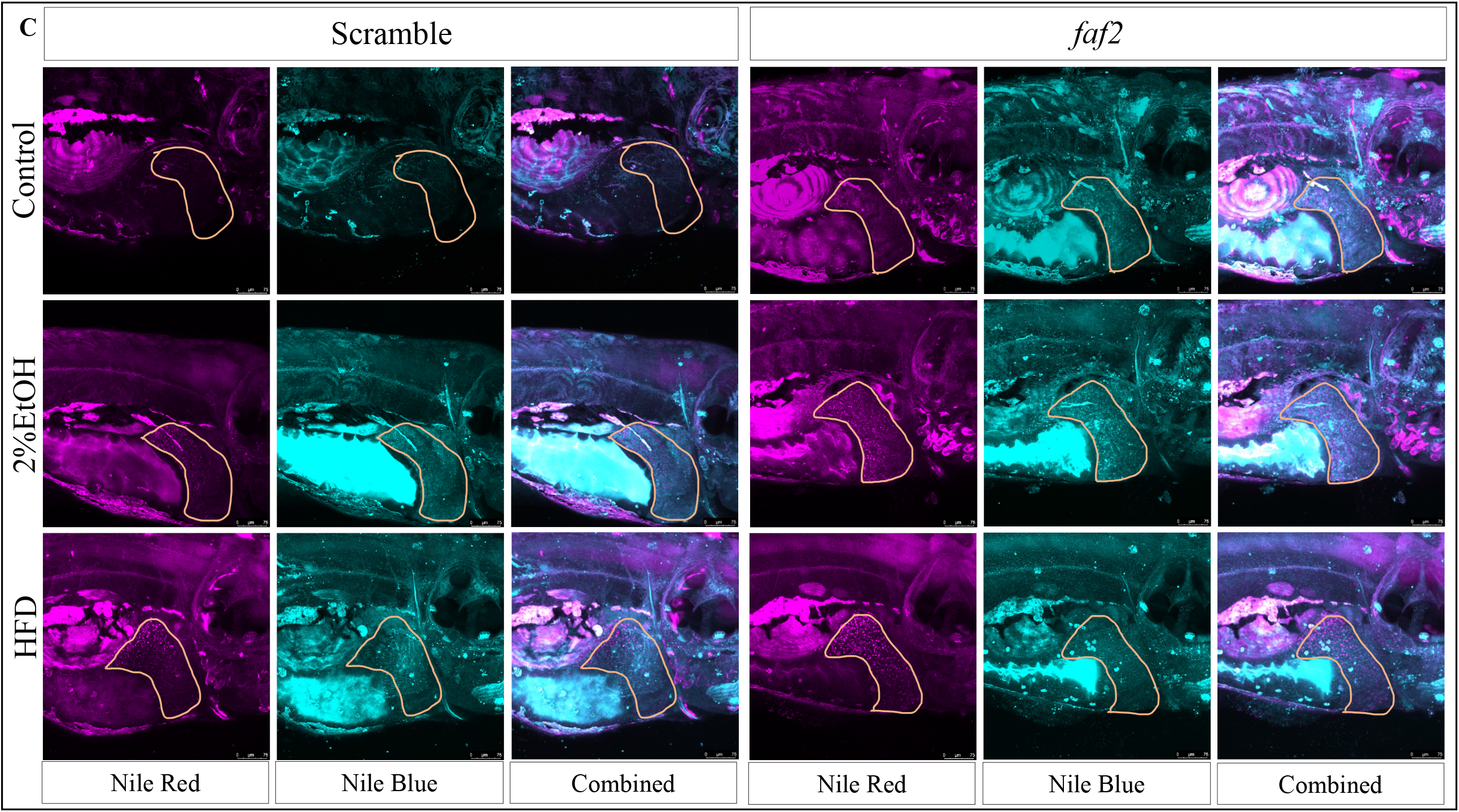
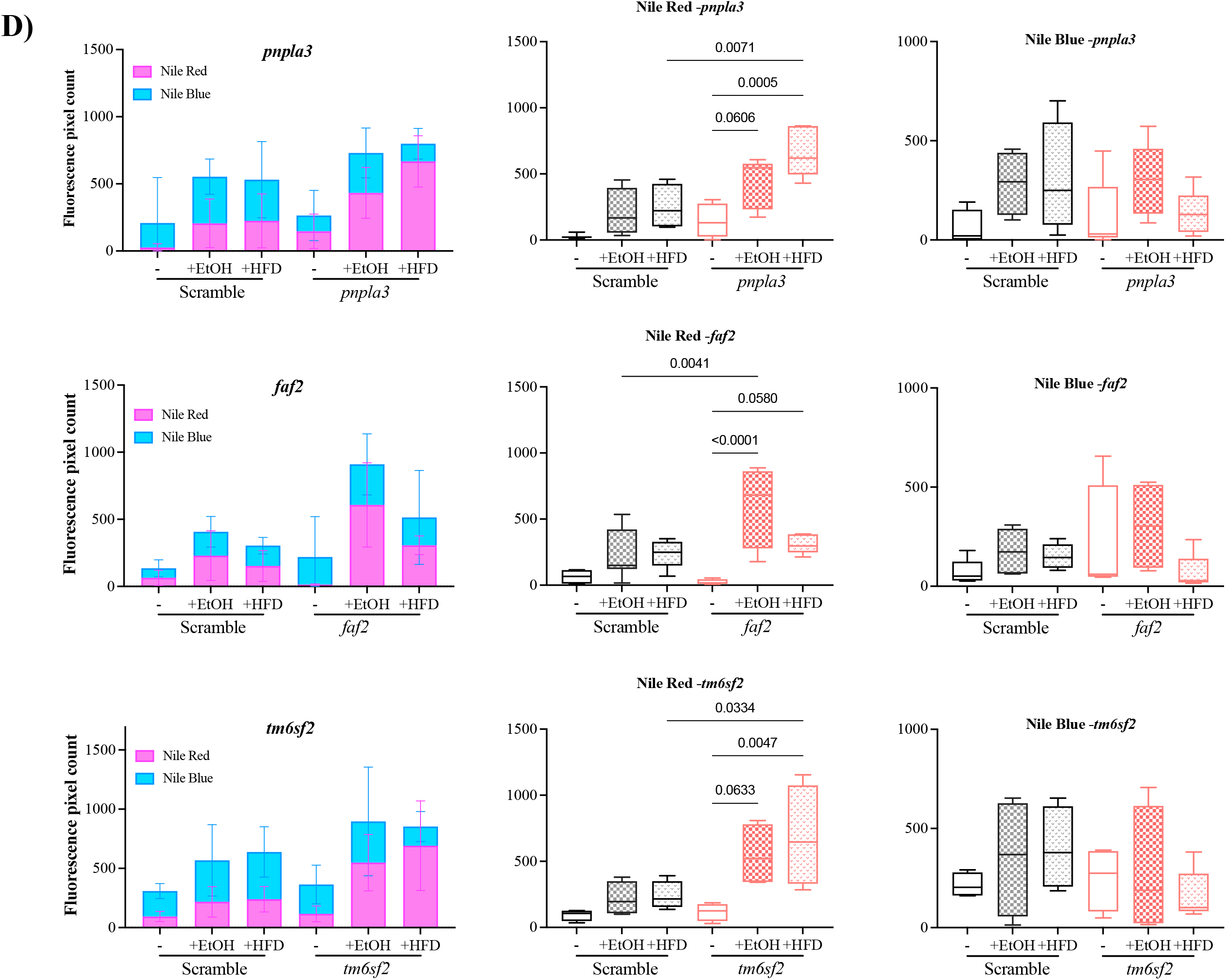
Ethanol or HFD exposure altered expression of genes in lipid biology pathways in zebrafish larvae. Lipid biology genes expression in *pnpla3, faf2* and *tm6sf2* crispants on 5 dpf following 48 hours of exposure to **A)** 2% EtOH or **B)** HFD. qPCR analysis was used to detect the expression of lipid-related genes relative to 18S. The average fold change was normalized to the expression in scramble control. Y- axis represents the foldchange over scramble control (marked in dotted line). Each point represents data pooled from10–15 zebrafish larvae per treatment. Error bars indicate interquartile range of the median. The ordinary one-way ANOVA with selected pair test was used to calculate the statistical significance.

Other genes in lipid biology, *hmgcra* (metabolism), *fabp10a* (transport), *notch1* (signalling), *serpb1* (synthesis) and *echs1* (β-oxidation) were increased by EtOH or HFD exposure in crispants but were not affected by knocking down *pnpla3, faf2* or *tm6sf2* alone.

### Live imaging with Nile Red and Nile Blue showed increased hepatic lipid droplets in crispant models

Live imaging of larvae was used to detect the hepatic localisation of neutral lipids, stained with Nile Red, in *pnpla3, faf2 and tm6sf2* crispants compared to their respective controls in the presence of EtOH and HFD (Figures 4A-C). Quantification demonstrated a significant increase of hepatic Nile Red staining in *pnpla3* crispants exposed to HFD, *faf2* crispants exposed to EtOH, and *tm6sf2* crispants exposed to HFD (Figure 4D).

**Figure 4:**
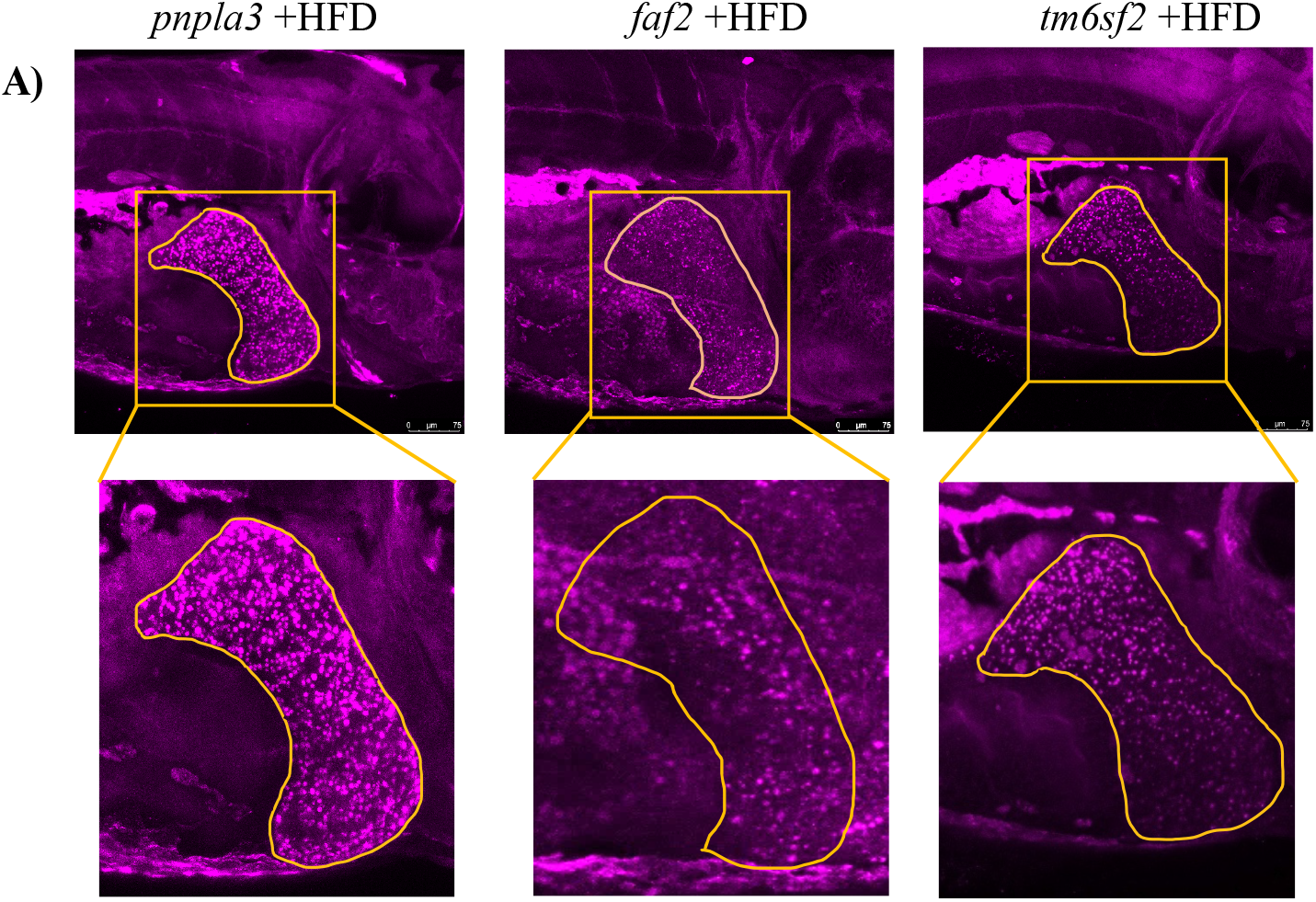
Live imaging of Nile Red and Nile Blue staining in *pnpla3, faf2 and tm6sf2* crispants treated with Ethanol/HFD exposure developed lipid droplets and free fatty acids. Live images show crispants **A)** *pnpla3* **B)** *faf2 and* **C**) *tm6sf2* treated with 2% EtOH or HFD and stained with Nile Red (magenta) and Nile Blue (cyan) using confocal microscopy. A z-stack of 75 µm was compiled for each sample. Highlighted region (orange line) is the liver of each zebrafish larvae. Images were used to quantify lipid (magenta fluorescence intensity), and free fatty acid (cyan fluorescence intensity) content using Fiji ImageJ software. Bar chart **D)** shows the amount of lipid and free fatty acids florescence count in each crispants. A set of 4-6 larvaewere used for the image analysis.

Conversely, Nile Blue staining, typically binding unsaturated free fatty acids, was unchanged in crispants versus scramble controls across treatment groups (Figure 4D). Quantification of total lipid staining area confirmed a significant increase in total lipids compared to scramble controls, with a major proportion of neutral lipids (Nile Red) compared to free fatty acids (Nile Blue) in all three crispants treated with EtOH or HFD (Figure 4D). Overlaying images showed no overlap between Nile Red and Nile Blue staining in the liver (Figure 4).

We also observed that the percentage of larger sized lipid droplets (>50 *μ*m^2^ vs 1-10 *μ*m^2^ or 11-50 *μ*m^2^) increased log10-fold in the livers of all three (*pnpla3, faf2, tm6sf2*) models compared to scramble controls (Figure 5A & B).

**Figure 5:**
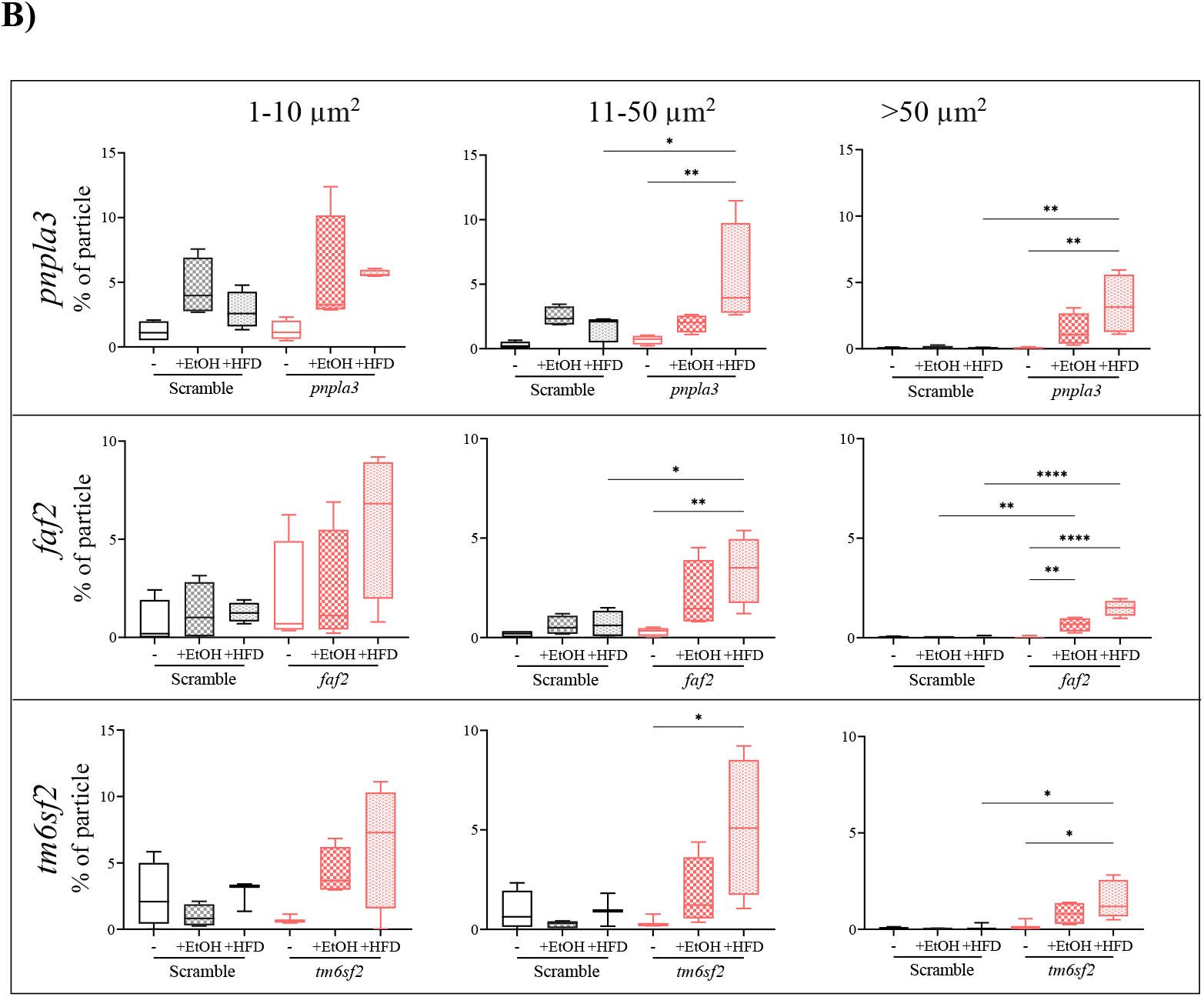
Nile Red staining showed larger size lipid droplets in 2%Ethanol or HFD treated crispants. **A)** Representative images show the liver area selected to calculate the lipid droplets size count in *pnpla3, faf2 and tm6sf2* crispants treated with HFD, stained with Nile Red (magenta) and imaged using confocal microscopy. **B)** Percentage of number of lipid droplet size (µm^2^), 1-10, 11-50 and greater 50 was measured using Fiji ImageJ software in scramble control and *pnpla3, faf2 and tm6sf2* larvae on 5 dpf following 48 hours of exposure to 2% EtOH and HFD.

## Discussion

We have used knockdown of *pnpla3, tm6sf2*, and *faf2* in zebrafish larvae to provide evidence of increased liver injury (lipid accumulation and neutrophil infiltration) in the presence of EtOH and HFD as models of diet induced ALD and NAFLD. Knockdown of liver disease susceptibility genes and exposure to EtOH or HFD significantly altered the expression of genes involved in lipid synthesis, oxidation, and transport, manifesting in increased neutral lipid accumulation. Lipid droplet numbers and size were increased in the livers of all three crispants exposed to EtOH/HFD. Increased lipid droplet size is one of the known markers of liver injury in the pathogenesis of ALD and NAFLD.[30, 31] Our data suggests that the underlying mechanism by which these gene variants interact with alcohol and HFD may be related to governing the size of lipid droplets, in addition to the absolute amount of lipid accumulated. We also provide evidence for the first time that *faf2*, our novel meta-GWAS discovery, may also be mediating the pathogenesis of ALD through lipid-droplet regulation in these diseases.

*Interaction of gene knockdown with EtOH and HFD*

We used Nile Red which selectively stains natural lipids (TG, cholesterol) and Nile Blue staining free fatty acids to visualise and differentiate the different types of lipids that accumulated in our models.[29, 32] Knockdown of our candidate genes specifically increased hepatic neutral lipids compared to free fatty acids in the presence of both EtOH and HFD, suggesting that these three genes regulate triglyceride metabolism in ALD and NAFLD models.

Lipid accumulation in the liver can occur due to any or all of the following: excessive de novo lipogenesis, namely increased fatty acid and TG synthesis, reduced β-oxidation of fatty acids, and defective lipid transport out of the liver.[33] Our fatty liver models combined with *pnpla3, faf2*, and *tm6sf2* knockdown inhibited the expression of genes such as *atgl* (TG hydrolysis), *acox1* (β-oxidation) and induced *srebp1* (lipid synthesis), *fabp10a* (lipid transport) and *hmgcra* (lipid metabolism) which was coincident with the observed excess fat build-up in the liver. We hypothesise these observations are linked through the reduction of triglyceride breakdown and increasing triglyceride synthesis. Furthermore, we hypothesise these nodes of lipid metabolism may be targets for treatments.

Expression of alcohol and lipid metabolism-related genes (*cyp2y3, cyp3a65, hmgcra, hmgcrb, fasn, fabp10α, fads2, echs1)* are shown to be dysregulated in zebrafish models of fatty liver.[33, 34] Upregulation of Cytochrome P450 family 2, subfamily Y, polypeptide 3 (*cyp2y3* zebrafish ortholog of human *cyp2e1* induced by alcohol) accelerates the speed of EtOH metabolism and accumulation of acetaldehyde increasing liver damage[34]. Upregulation of *SREBP1c* transcription factor at the transcriptional level appears to be mediated by acetaldehyde, the toxic metabolite of alcohol metabolism.[35, 36] We hypothesise that increased *srebp1* expression is regulated by alcohol metabolite acetaldehyde in our zebrafish model of EtOH fatty liver, leading to TG accumulation and inflammation shown by increased neutrophil infiltration in the livers of all three knockdown disease models. However, the possibility of alternate mechanisms for *srebp1* increase, such as ER stress response cannot be ruled out in with HFD exposed crispants. The SREBP transcription factors (SREBP1, SREBP2) are well-characterized regulators of hepatic lipogenesis; function by increasing the expression of the other genes (*echs1*)[28] required for TG biogenesis.[37] Similar to our data in zebrafish, increased *de novo* lipogenesis in mice exposed to ethanol or HFD is thought to be through increased expression of SREBP1 in the liver.[3] Increase in HMG-CoA reductases (*hmgcra*), a key enzyme in lipid metabolism regulating cholesterol biosynthesis genes, and fatty acid-binding protein 10a (*fabp10a*), an intracellular fatty acid-binding protein involved in intracellular lipid metabolism and fatty acid transport[34], shows our genetically modified zebrafish models of fatty liver have several genes dysregulated in lipid synthesis/metabolism pathways resulting in hepatic lipid accumulation. Interestingly, knocking down genes itself did not significantly affect the expression of these genes suggesting no direct effect of *pnpla3, faf2* and *tm6sf2*. However, EtOH or HFD exposure significantly upregulated expression of these genes in the crispants versus respective scramble controls, indicating *pnpla3, faf2 and tm6sf2*- dependent interactions with EtOH/HFD mediating injury through one or more (acetaldehyde, TG biogenesis, cholesterol biosynthesis) pathways in these models.

*Role of pnpla3, faf2 and tm6sf2 in ALD and NAFLD (Box 1)*

**Box 1:**
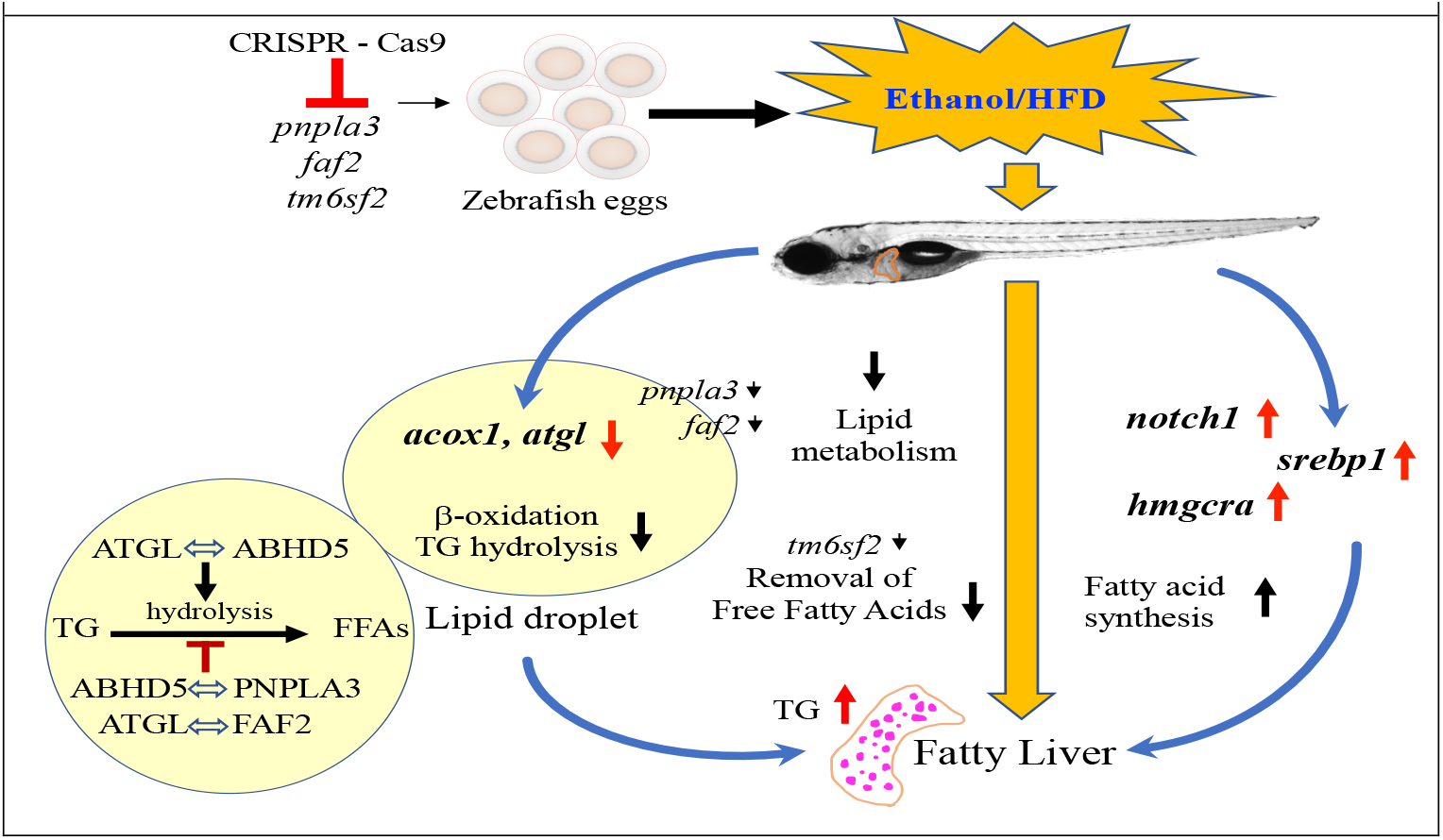
Model of *pnpla3, faf2 and tm6sf2 role in ALD and NAFLD:* Lipid metabolism genes/pathways (blue arrows); Dysregulation of genes in zebrafish models (red arrows); affected lipid processes (black arrows); Lipid droplets showing increased TG (magenta) in the zebrafish larva liver.

In our *pnpla3* model, knockdown itself increased hepatic lipid accumulation which further increased in the presence of EtOH and HFD. This increase could result from reduced expression of *atgl* and *acox1* involved in TG hydrolysis and β-oxidation respectively, and increased expression of *echs1* and lipid synthesis genes *srebp1* and *hmgcra*. PNPLA3 proteins reside on lipid droplets and bind to α/β hydrolase domain 5 (ABHD5) thereby suppressing ABHD5 binding to adipose triglyceride lipase (ATGL), the major TG hydrolysing enzyme,[38] and ABHD5/ATGL-dependent lipolysis.[39] We suggest that *pnpla3* in our model functions by increasing TG accumulation and lipid droplets through inhibiting ABHD5/ATGL lipolysis pathway (Box 1). We show evidence specifically of increased TG accumulation in the liver through Nile Red staining in *pnpla3* knockdown models and reduced *atgl* further in the presence of EtOH/HFD.

Interestingly, FAF2 inhibits lipid droplet degradation by binding to ATGL, also inhibiting the binding with ABHD5 described above and in Box 1. In our model, knocking down *faf2* decreased *atgl* expression, however, as there was no *ABHD5* ortholog available in zebrafish we could not check its expression. Similar to *pnpla3*, knockdown of *faf2* reduced expression of β-oxidation genes. Although, the *faf2* HFD model increased lipid synthesis (*srebp1* and *hmgcra)* and lipid signalling *(notch1)* significant accumulation of TG (Nile Red) was only evident in the presence of EtOH. So far, *FAF2* has not been reported to be associated with NAFLD[40] but only with ALD[7] but whether it is specific to ALD risk is yet to be determined. FAF2 is a sensor of intracellular levels of long-chain unsaturated fatty acids and is involved in TG metabolism[41], but the exact functions of FAF2 in lipid metabolism are unknown. We propose that FAF2 maintains lipid homeostasis by regulating TG hydrolysis in a self-organising system when activated by excess free fatty acids dissociating ATGL from ABHD5 thereby inhibiting TG hydrolysis in lipid droplets. In light of our *faf2* knockdown model increasing TG, and the reported association of *FAF2:*rs374702773 variant with reduced alcohol-related cirrhosis[7], it can be speculated that *FAF2:*rs374702773 mutation may be a gain of function. We also hypothesise the possibility of an interaction between PNPLA3 and FAF2 as they both are involved in inhibiting TG hydrolysis and regulating TG accumulation via the ATGL-ABHD5 pathway.

Loss of TM6SF2 function modulates hepatic lipid metabolism to produce significant changes in intracellular lipid accumulation.[42] Knockdown of *TM6SF*2 which assists secretion of lipids in TG-rich lipoprotein from lipid droplets increases intracellular lipid deposition and is accompanied by enhanced fatty acid synthesis and impaired fatty acid oxidation in mice.[8, 42] Our *tm6sf2* knockdown alone confirms this role by downregulating β-oxidation with increased TG accumulation, and further by increasing lipid synthesis in the presence of EtOH and HFD.

### Strengths and limitations

The strength of the study lies in the use of zebrafish as genetic models for studying fatty liver diseases as liver development and function are well conserved with humans.[43] Our use of transparent zebrafish larvae allowed live imaging to visualise hepatic lipids *in situ* using Nile Red and Nile Blue staining. As we used a mosaic gene knockdown technique that reduced the gene expression instead of completely deleting the genomic locus, these findings could also be replicated in stable mutants/knockouts to confirm the proposed mechanisms. Inablility to investigate the genes for which human orthologs are not available in zebrafish is also a limitation of the study model and should be replicated in other experimental models.

In conclusion, we have created genetic and fatty liver disease models showing that *pnpla3, faf2* and *tm6sf2* are involved in lipid accumulation via dysregulated triglyceride metabolism in EtOH- and HFD-diet induced liver injury. Future studies are required using double knockouts to clarify gene x gene interactions (eg. *pnpla3, faf2*) for better understanding of mechanisms influencing these diseases and identify novel therapeutic molecules to target accumulation of lipid early for prevention of disease progression.

## Acknowledgment

We thank the Centenary imaging facility core and Sydney Cytometry staff Dr Kristina Jahn for their assistance.

