## Supplementary material for "Investigating the role of lipid genes in liver disease using fatty liver models of alcohol and high fat in zebrafish (*Danio rerio*)": Supplementary Document.docx

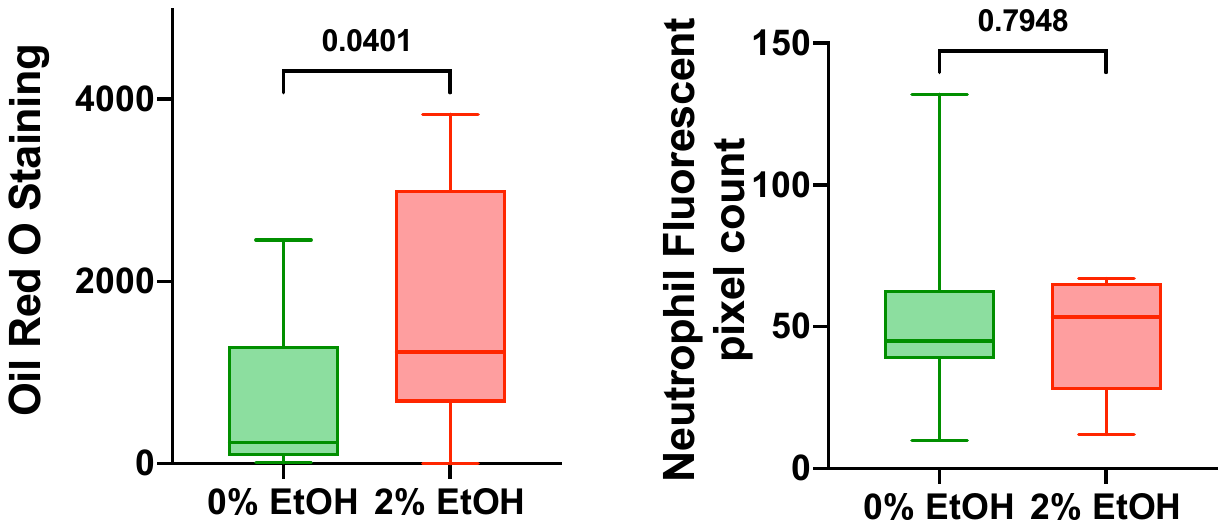

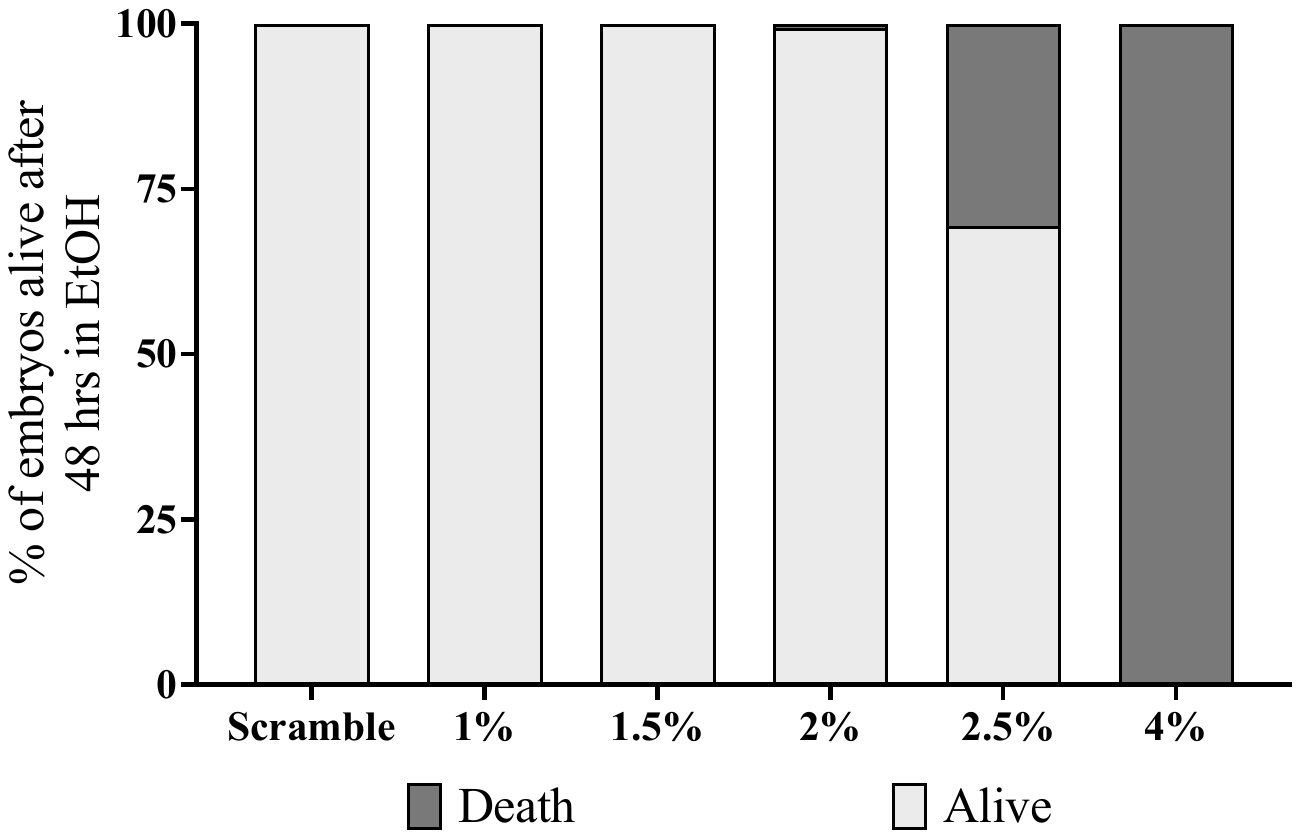

A)

C)

B)

0%EtOH

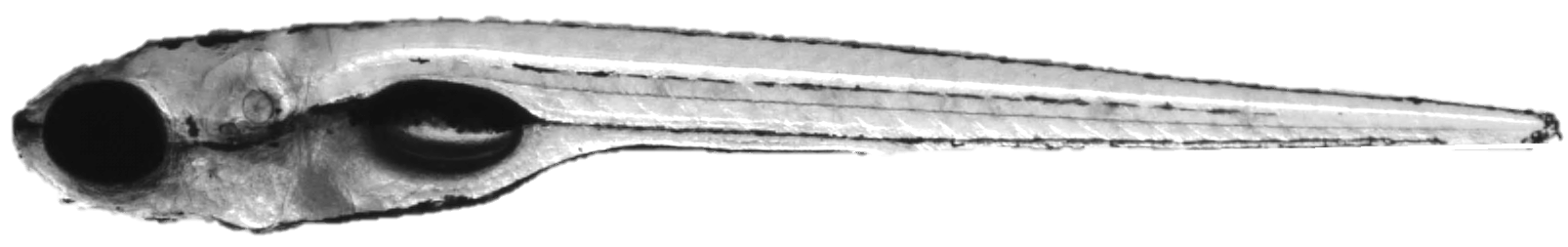

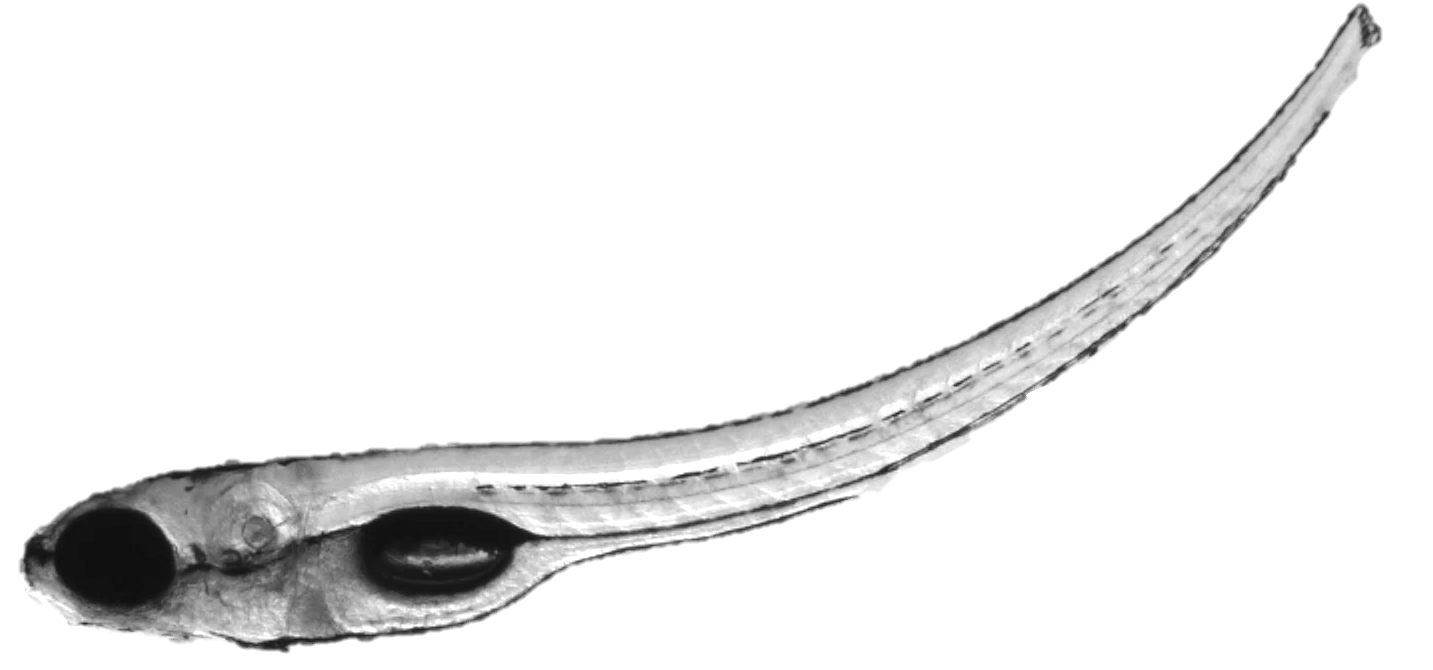

2%EtOH

% of larvae alive after 48 hrs in 2% EtOH

d)

**Supplementary Figure 1: Zebrafish larvae exposed to alcohol develop morphological abnormalities and steatosis** 5 dpf larvae treated with 0% or 2% EtOH for 48 hours. **A)** Mortality of zebrafish larvae in ethanol concentration **B)** The develop morphologic abnormalities including upward curvature of the trunk and tail. **C)** Quantification of lipid accumulation using wholemount oil red O staining for 48 hours reveals steatosis. **D)** Quantification of neutrophil shows no significant changes in liver.

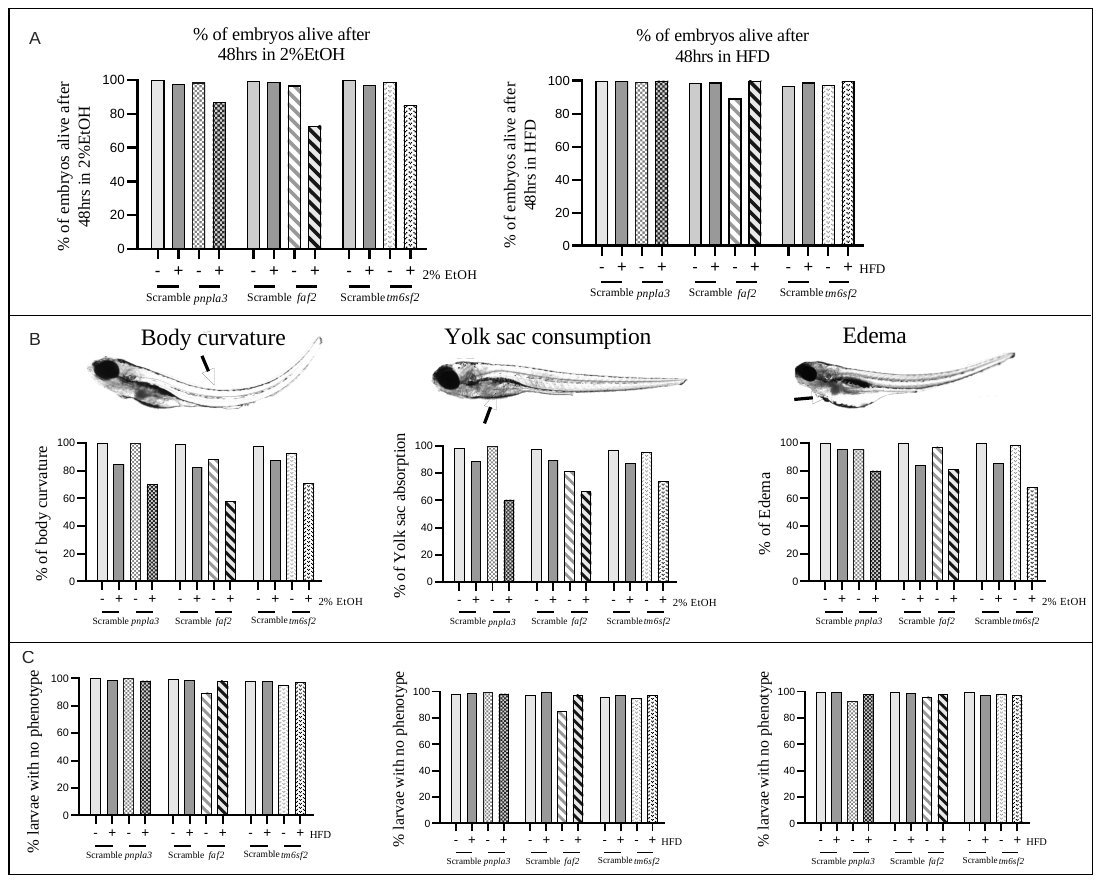

% of larvae alive after 48 hrs in 2% EtOH

% of larvae alive after 48 hrs in HFD

**Scramble**

***pnpla3***

***faf2***

Control

HFD

***tm6sf2***

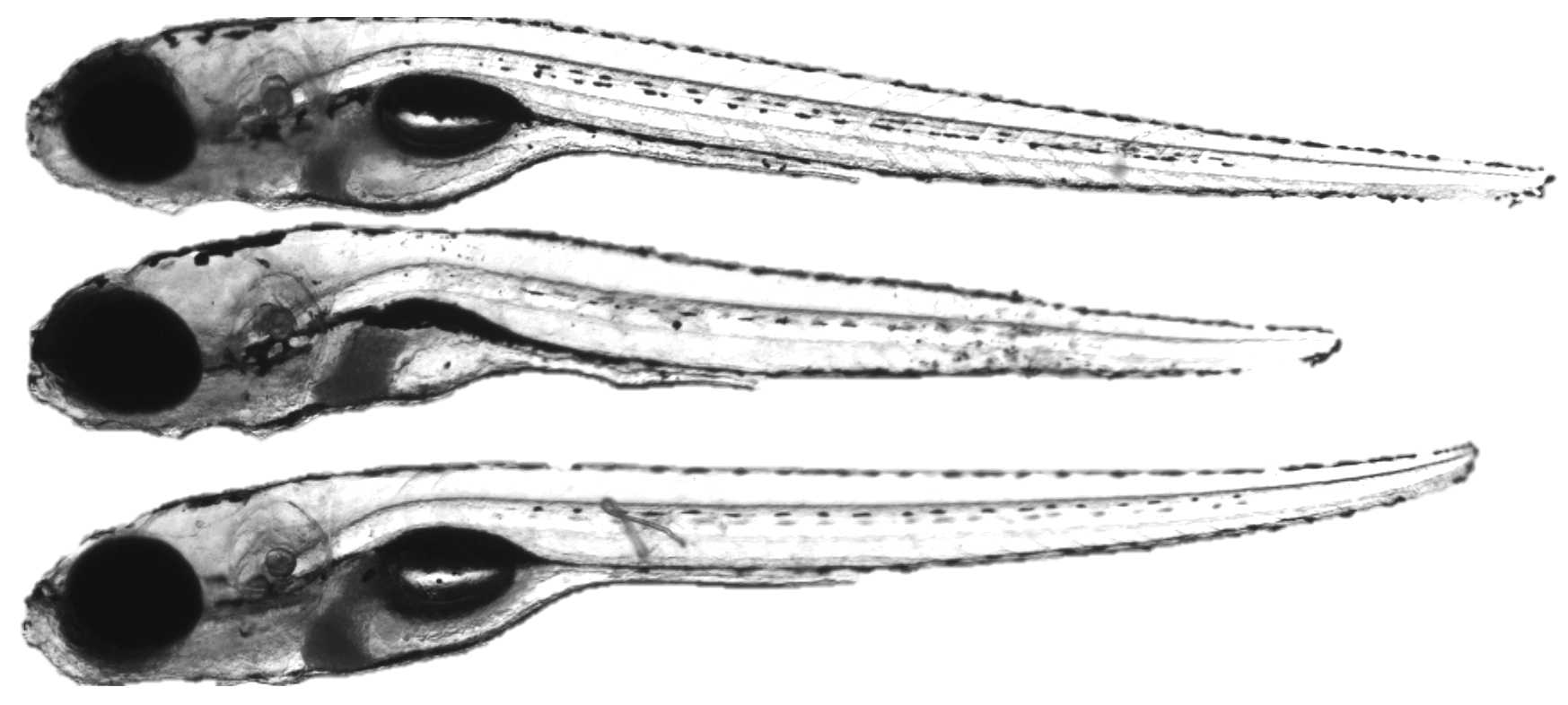

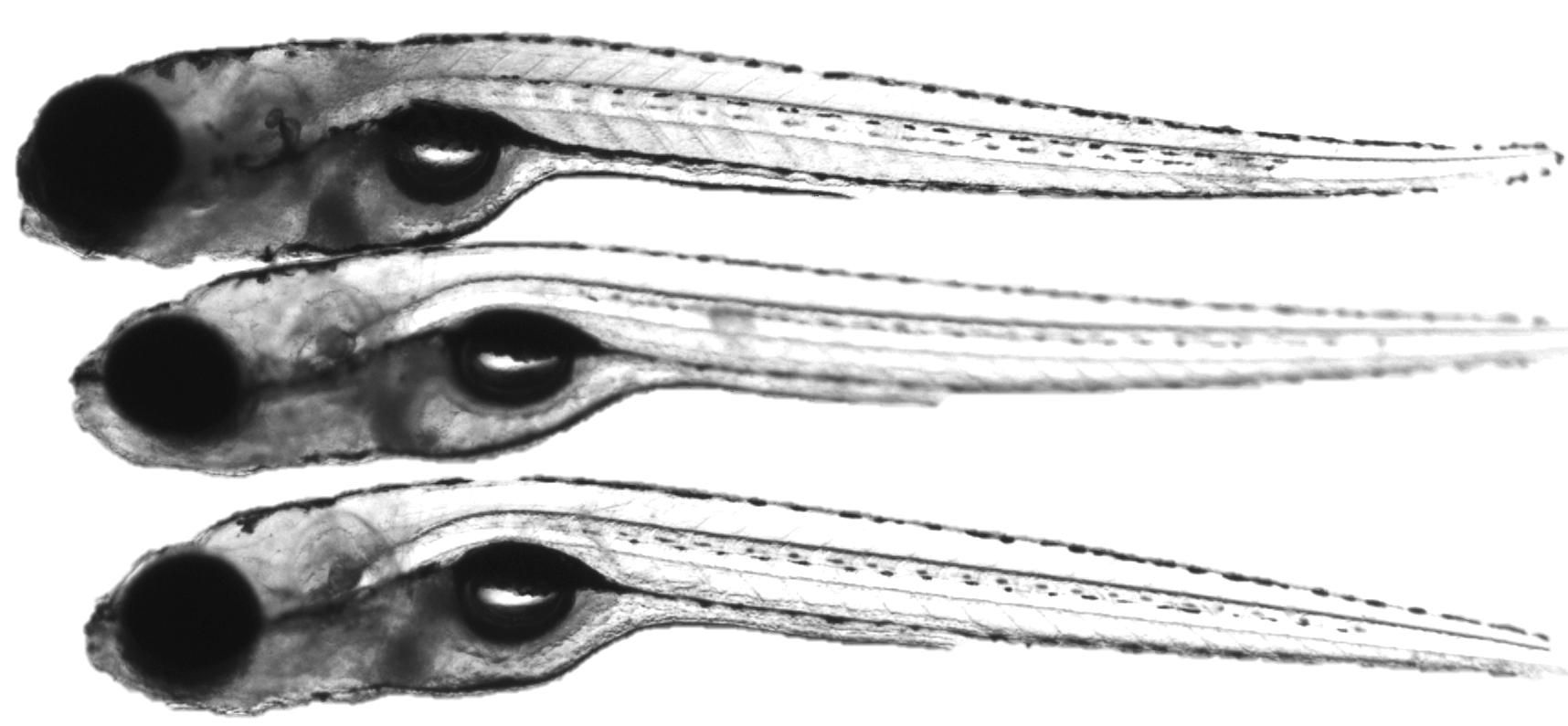

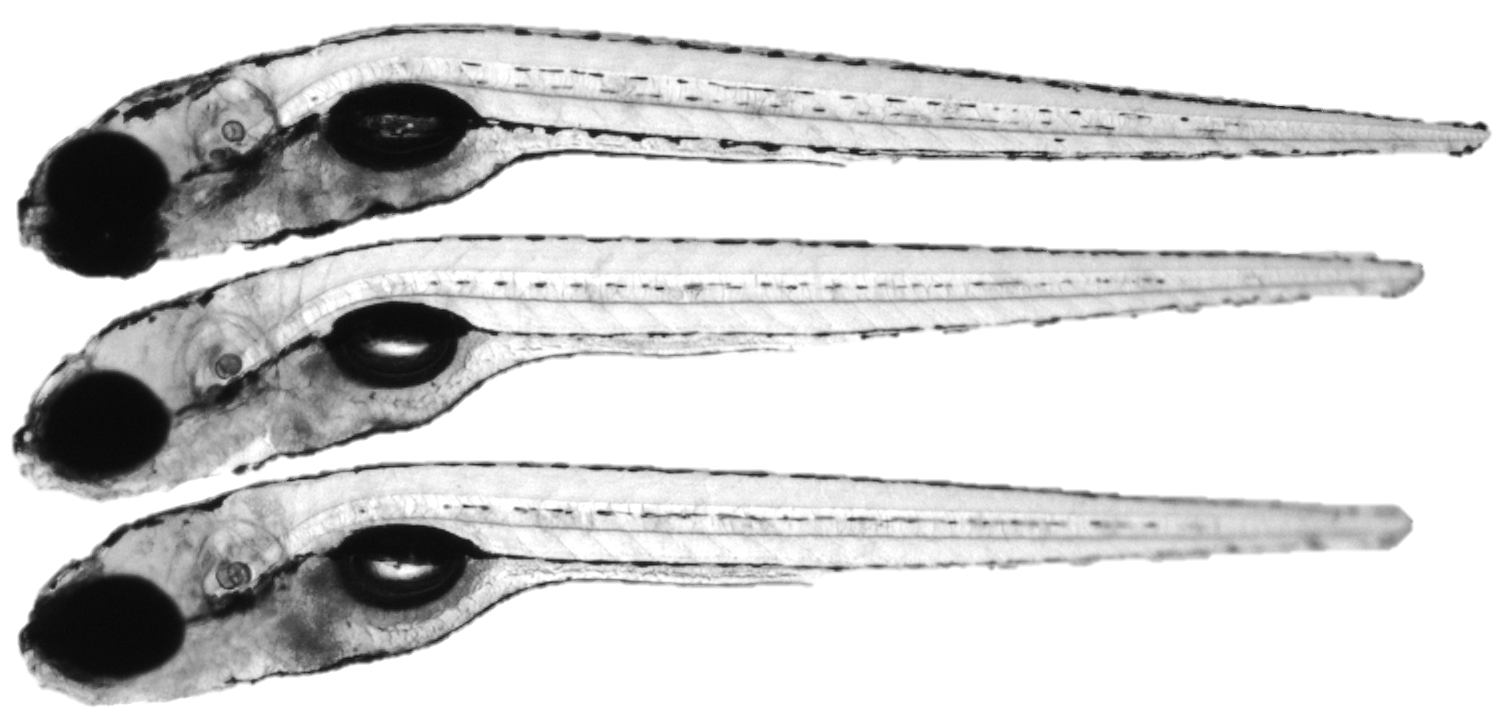

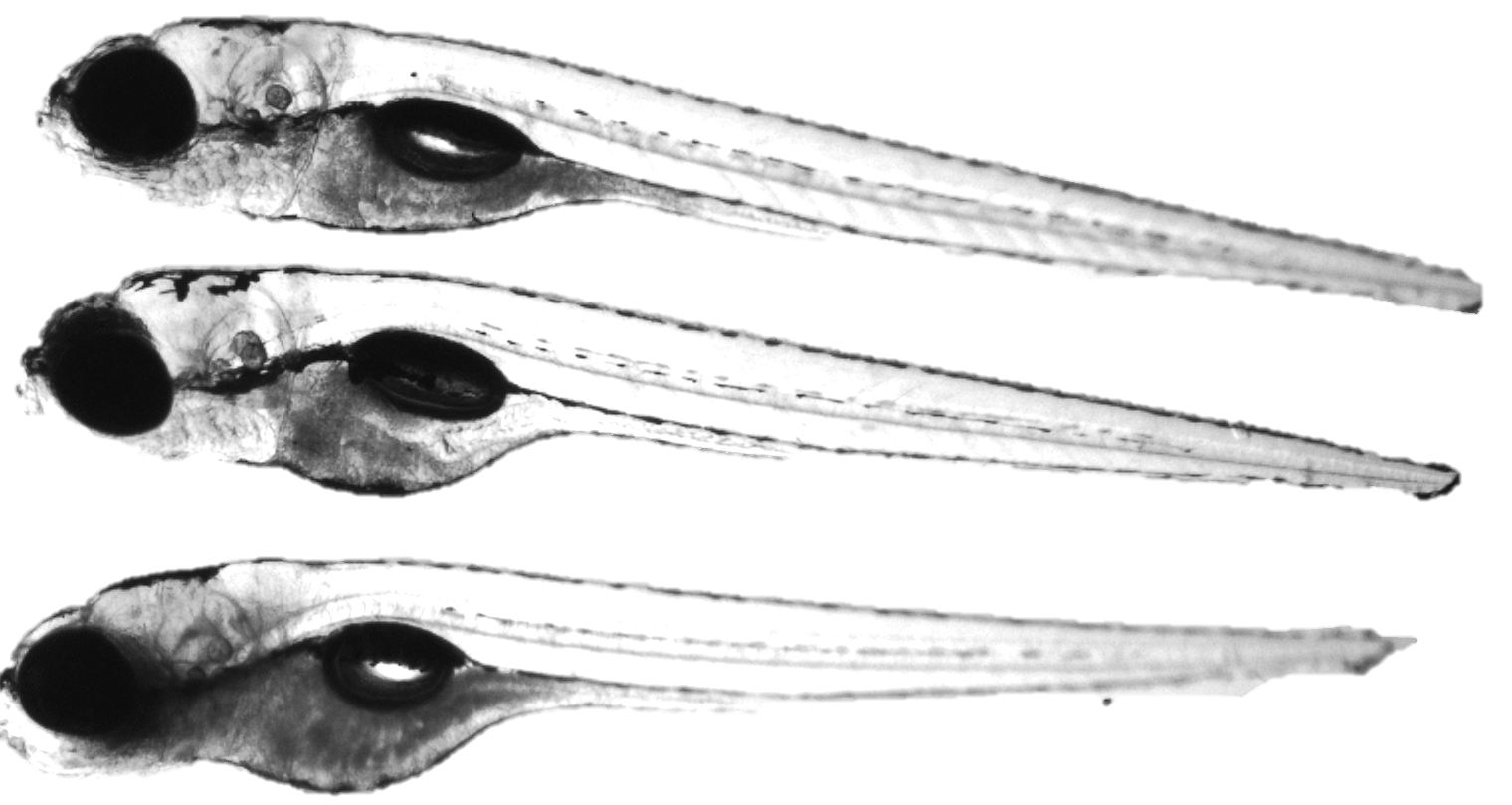

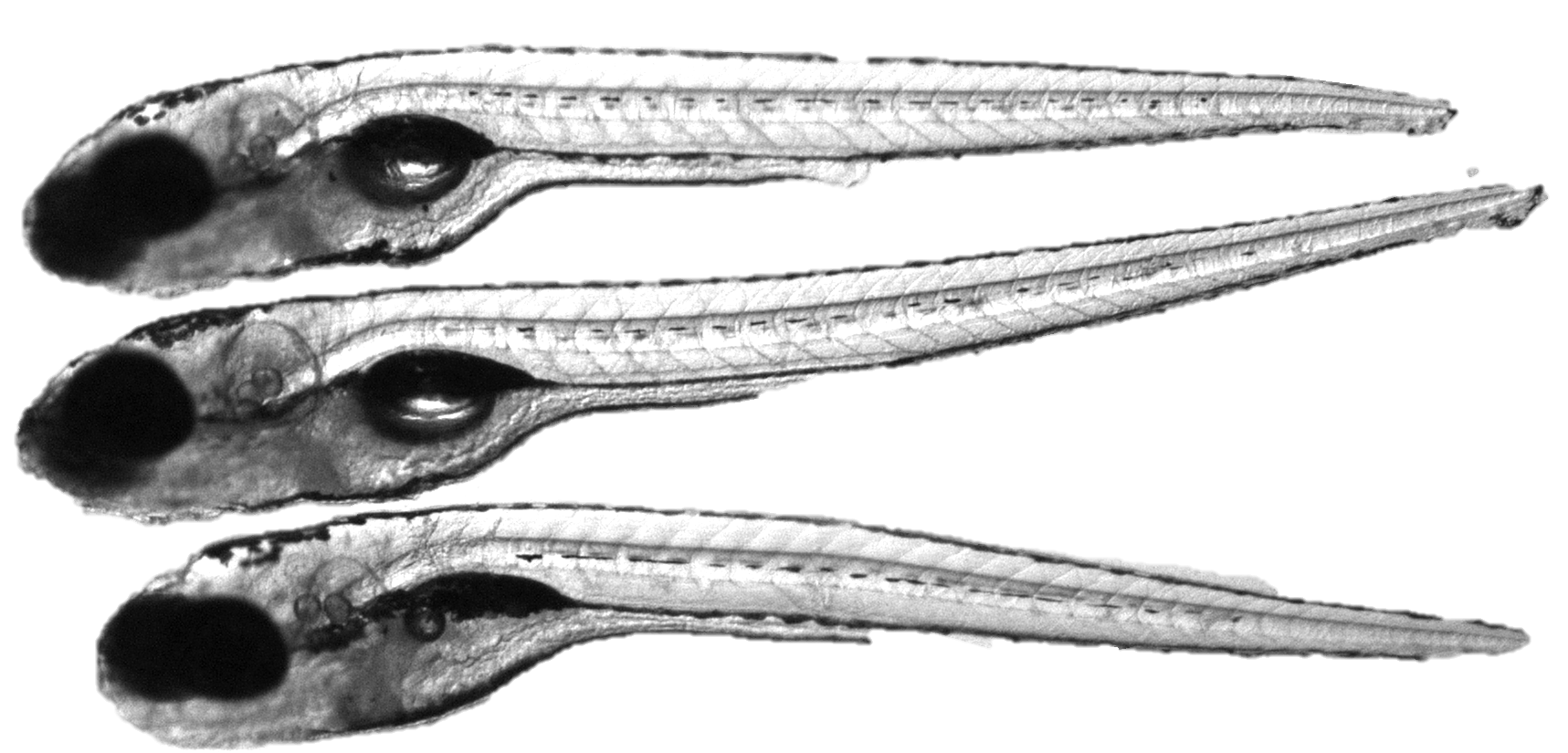

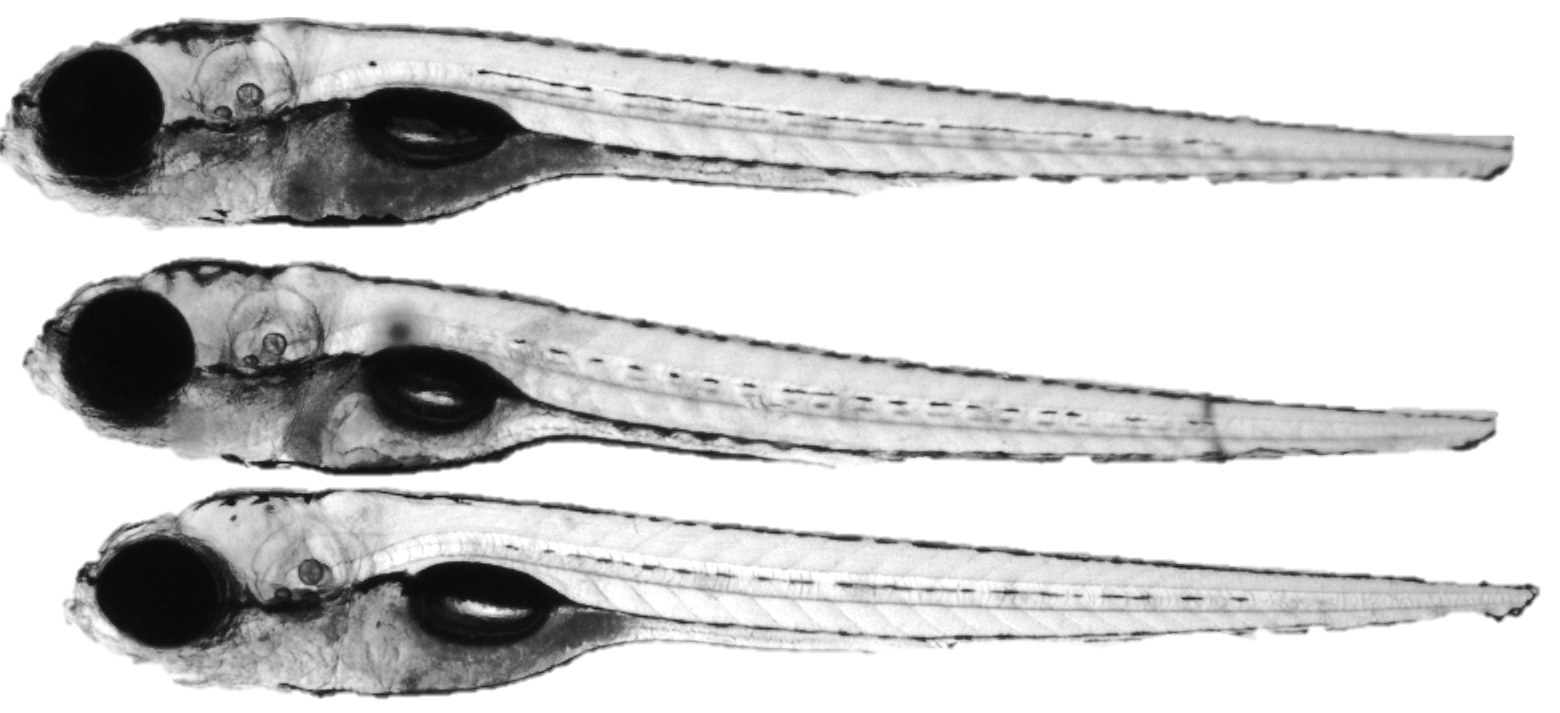

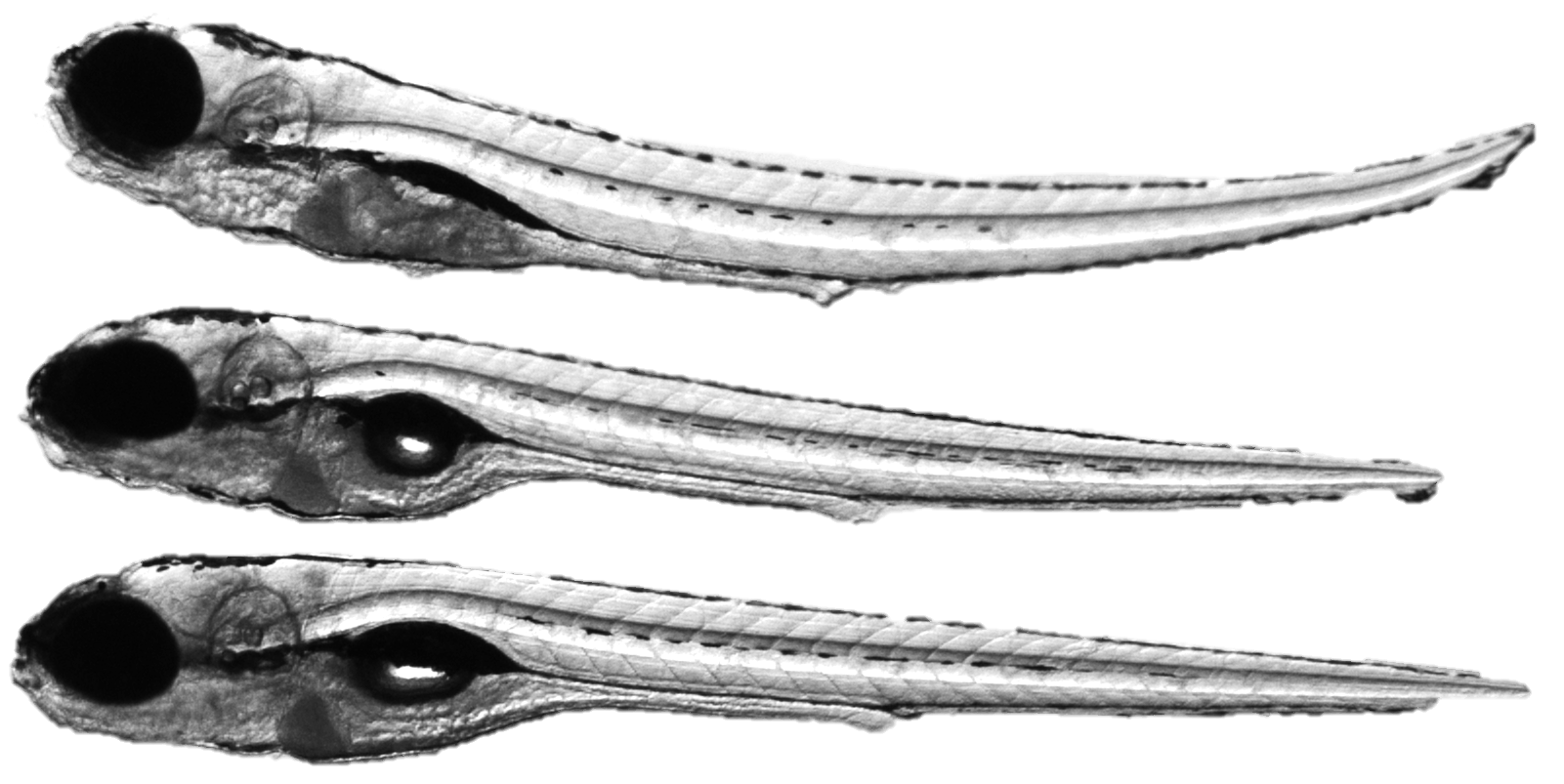

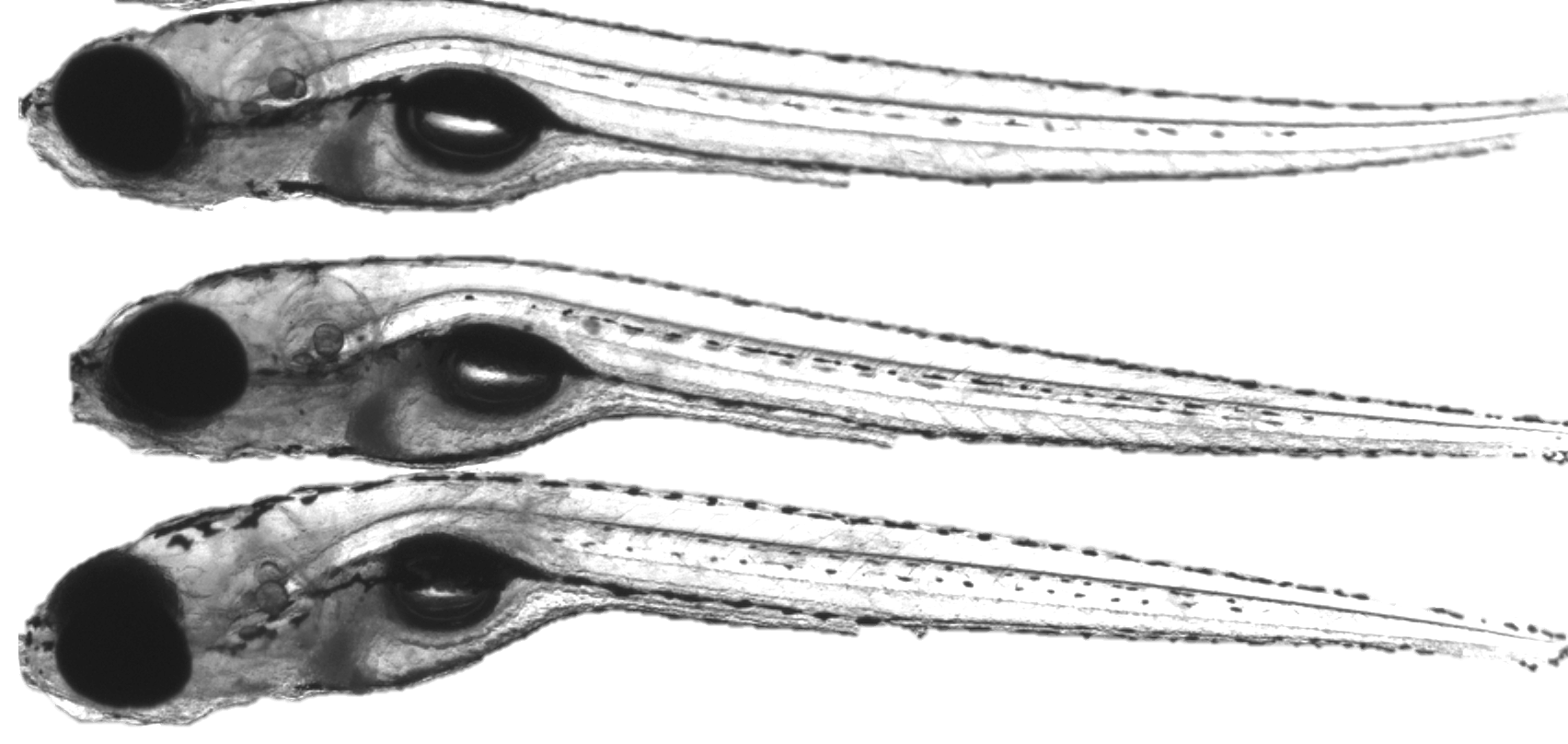

D)

**Supplementary Figure 2: Survival and morphological abnormalities in three gene crispants treated with EtOH or HFD for 48 hours. A)** mortality of crispant larvae after 48 hours. Percentage of morphological abnormalities in **B)** EtOH **C)** HFD exposed crispant larvae compared to controls. Experiments ware performed in three clutches and each clutch had 10-15 larvae per group. **D)** Brightfield images from live Tg(lyzc:egfp) showing a range of developmental morphological abnormalities in *pnpla3*, *faf2 and tm6sf2* crispants with HFD exposure.

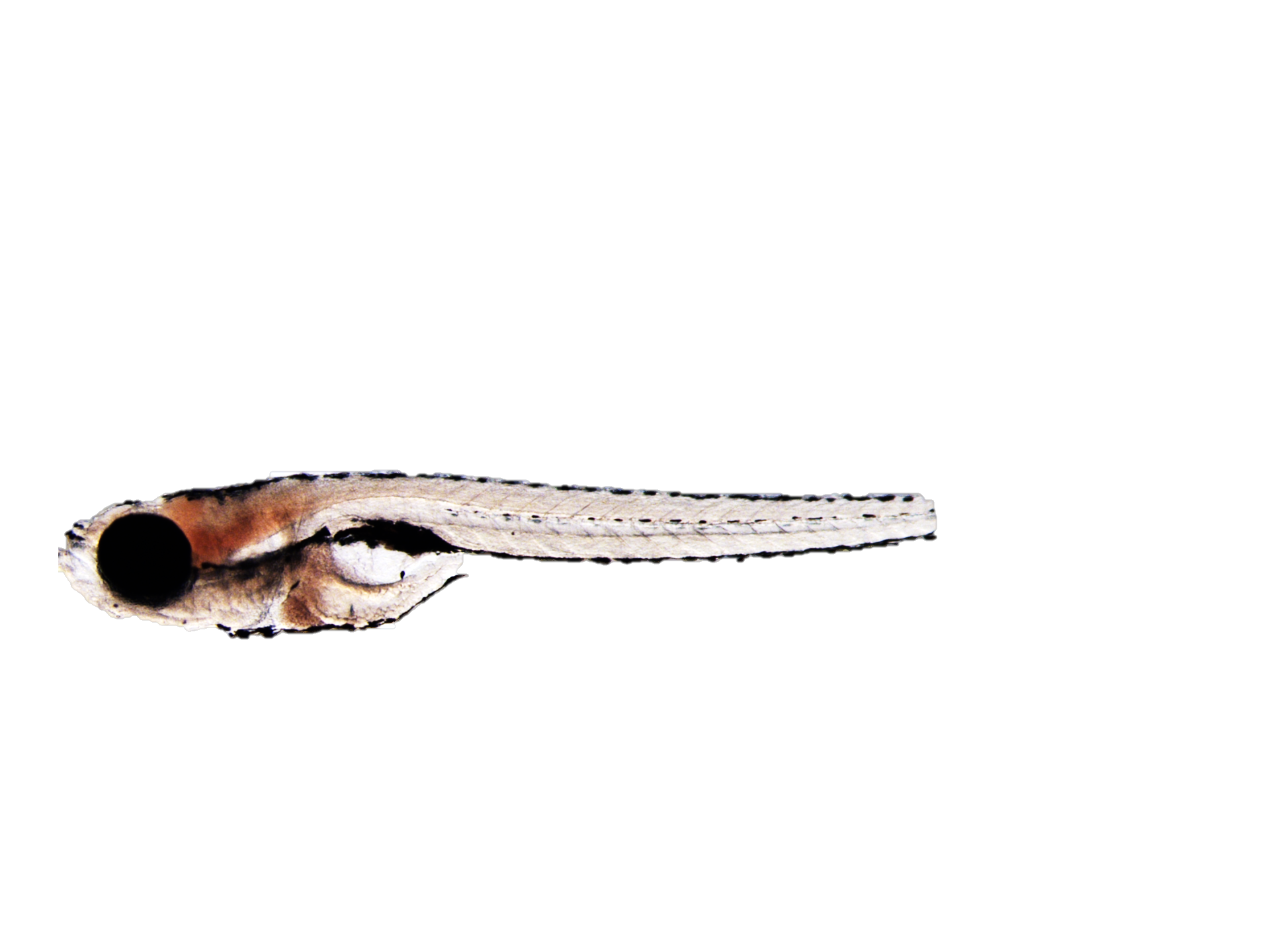

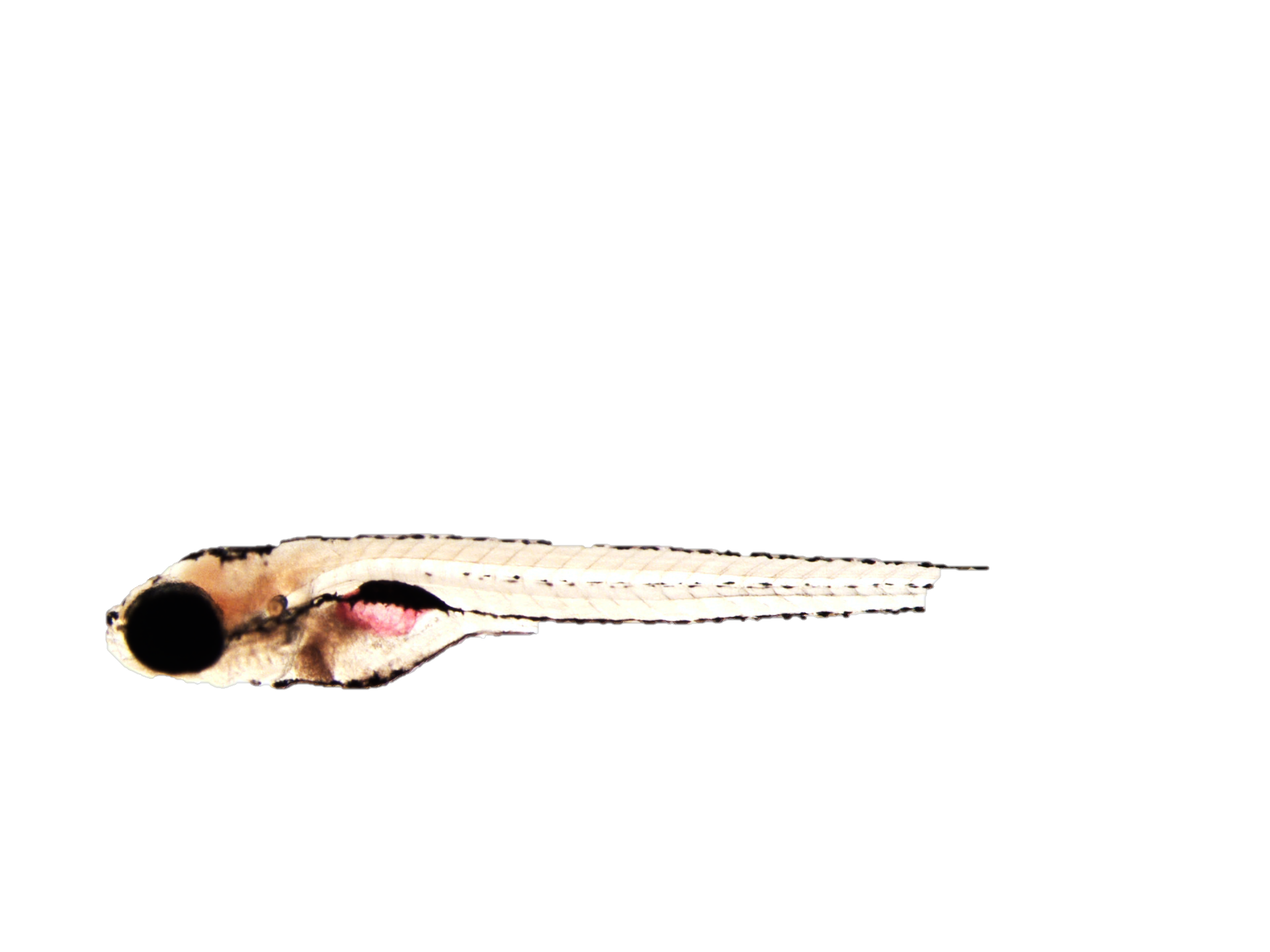

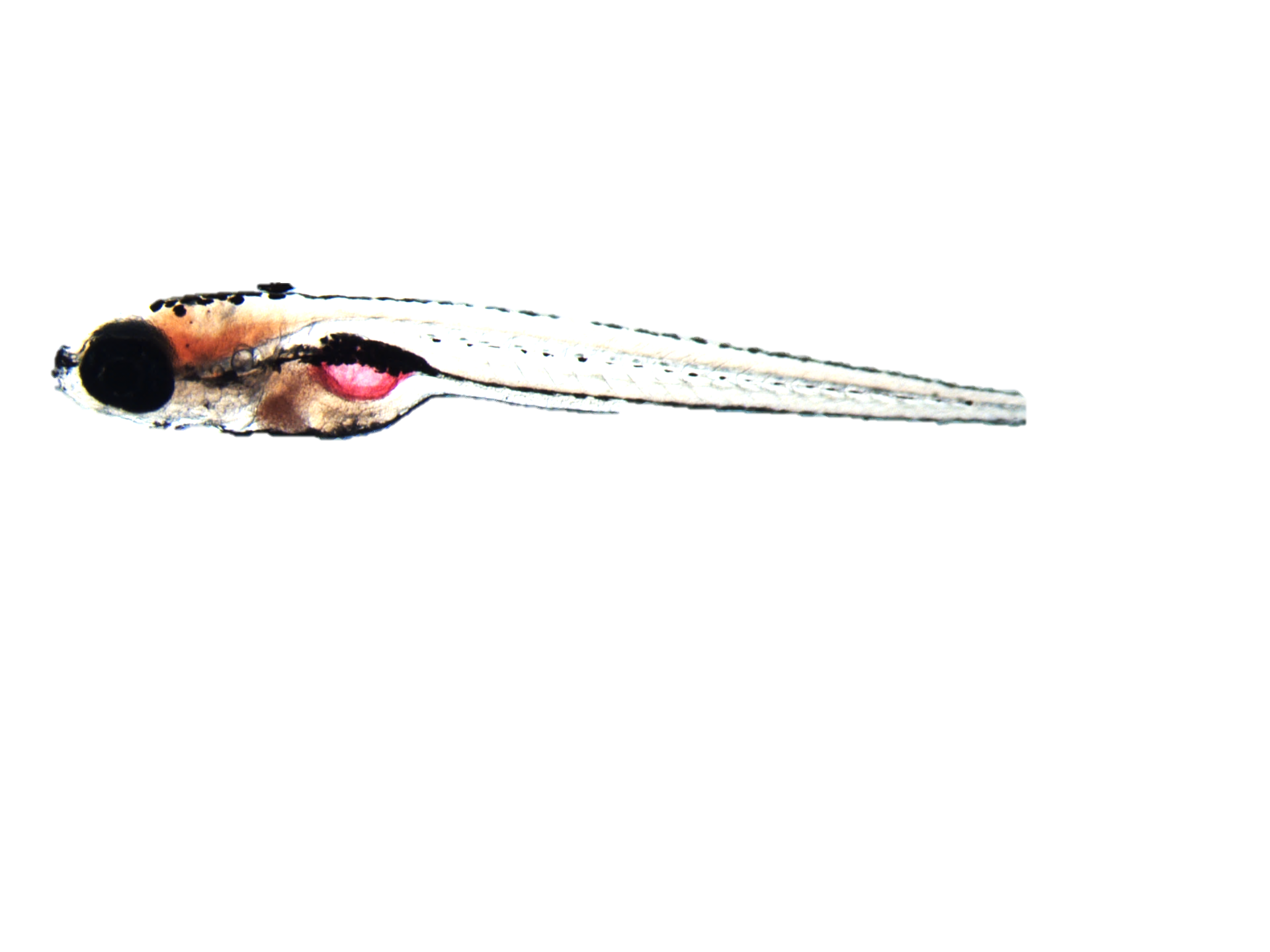

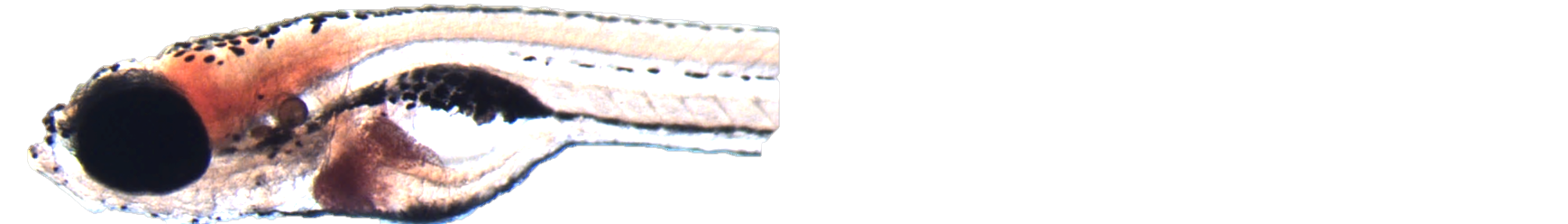

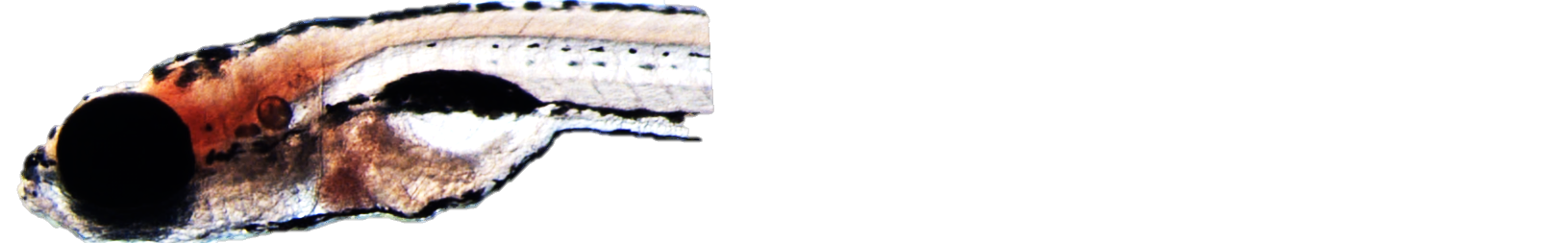

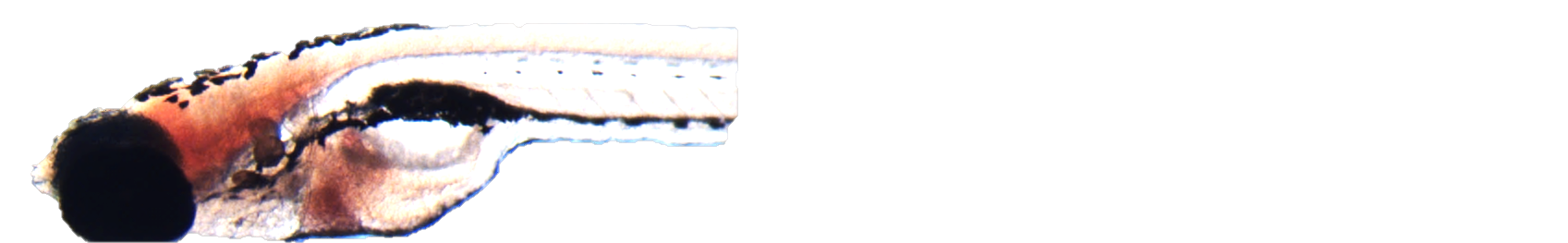

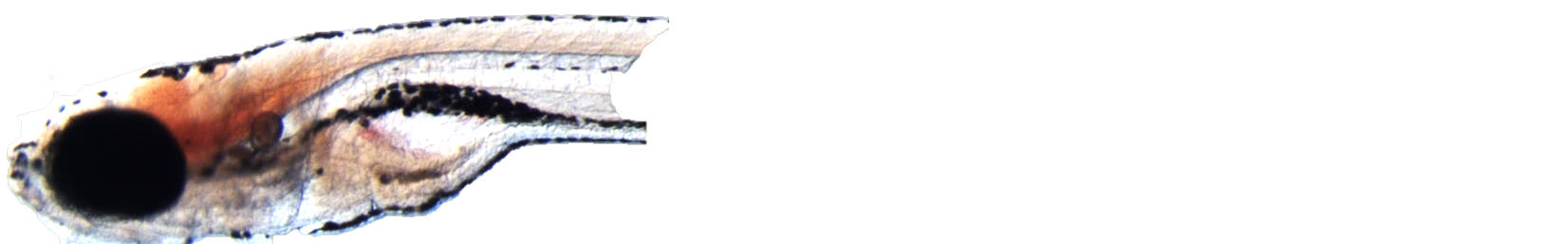

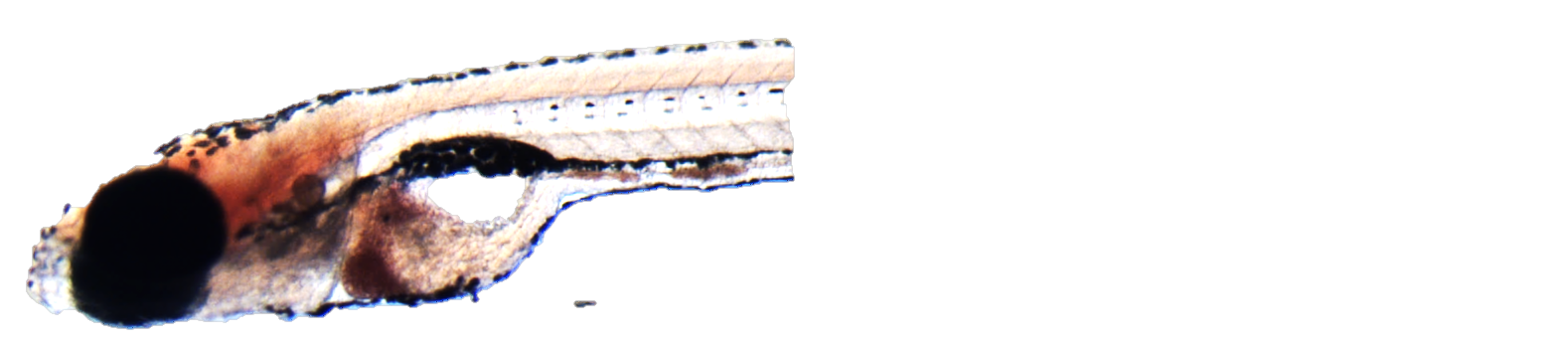

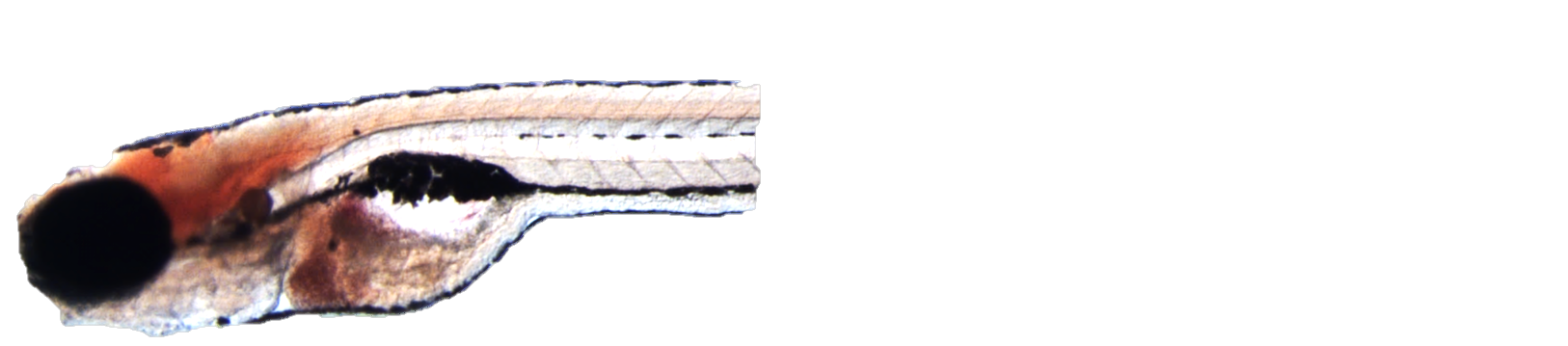

Control

*pnpla3*

2% EtOH

*faf2*

Scramble

HFD

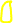

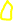

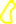

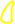

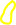

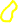

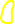

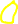

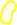

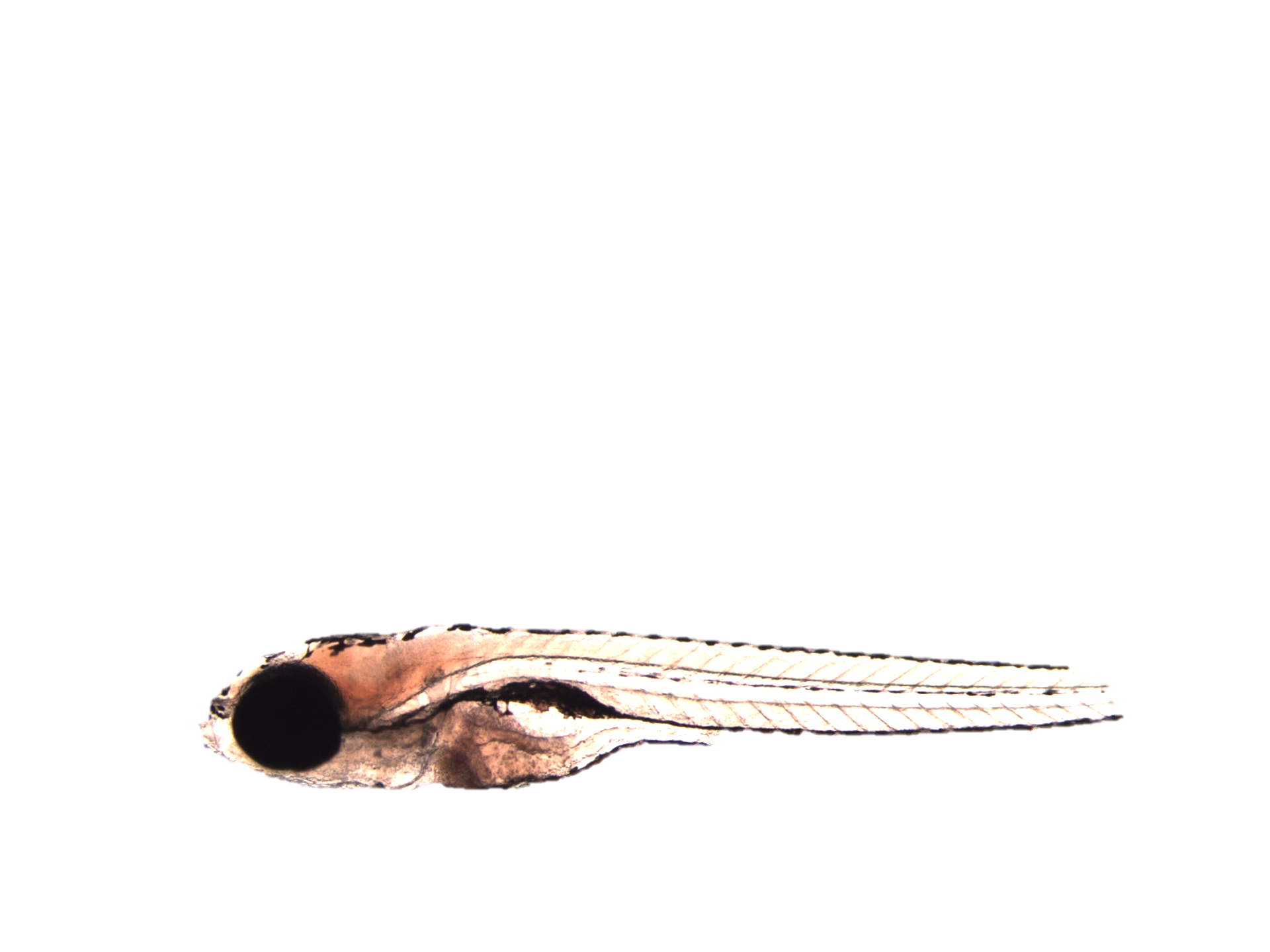

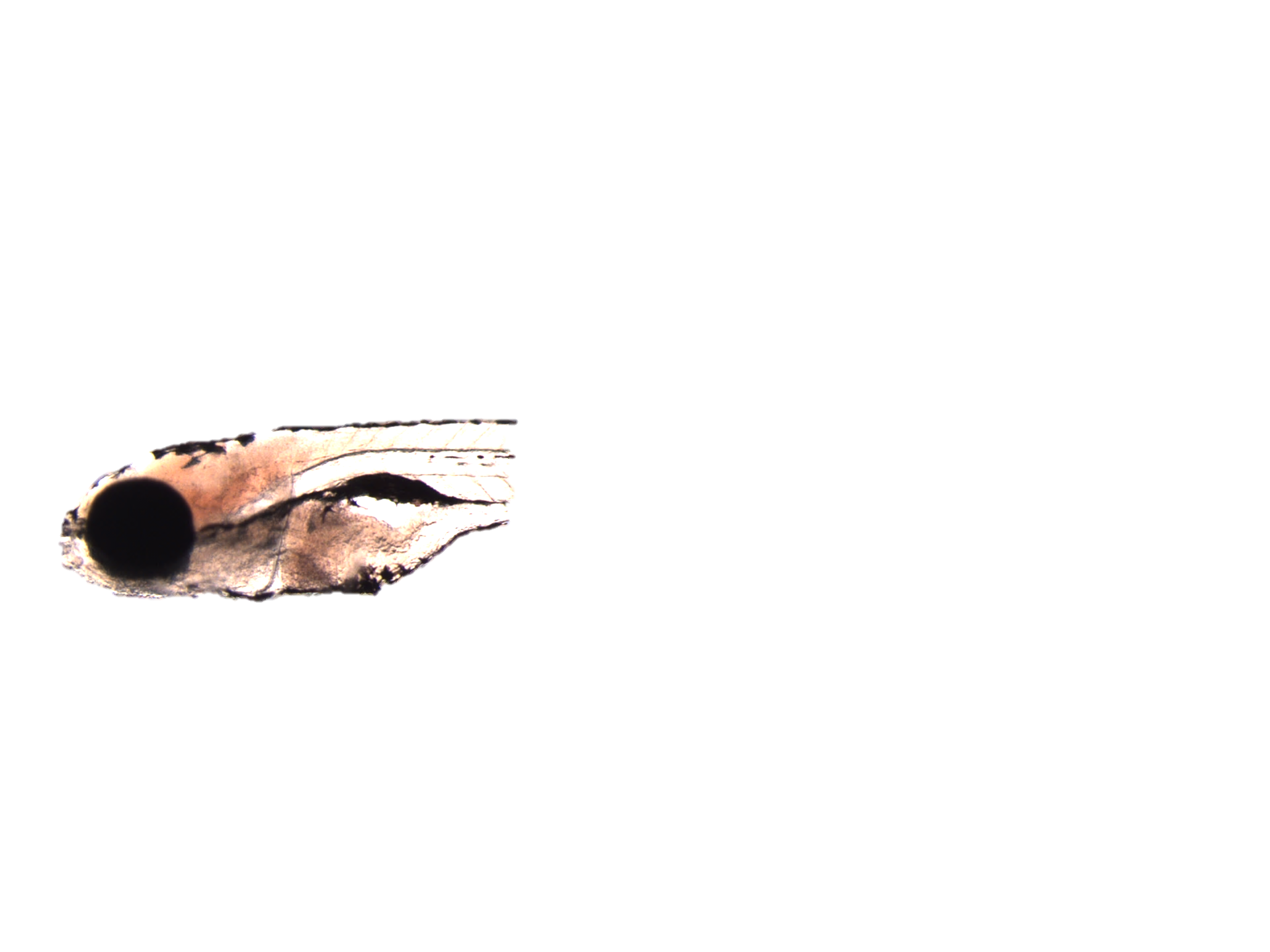

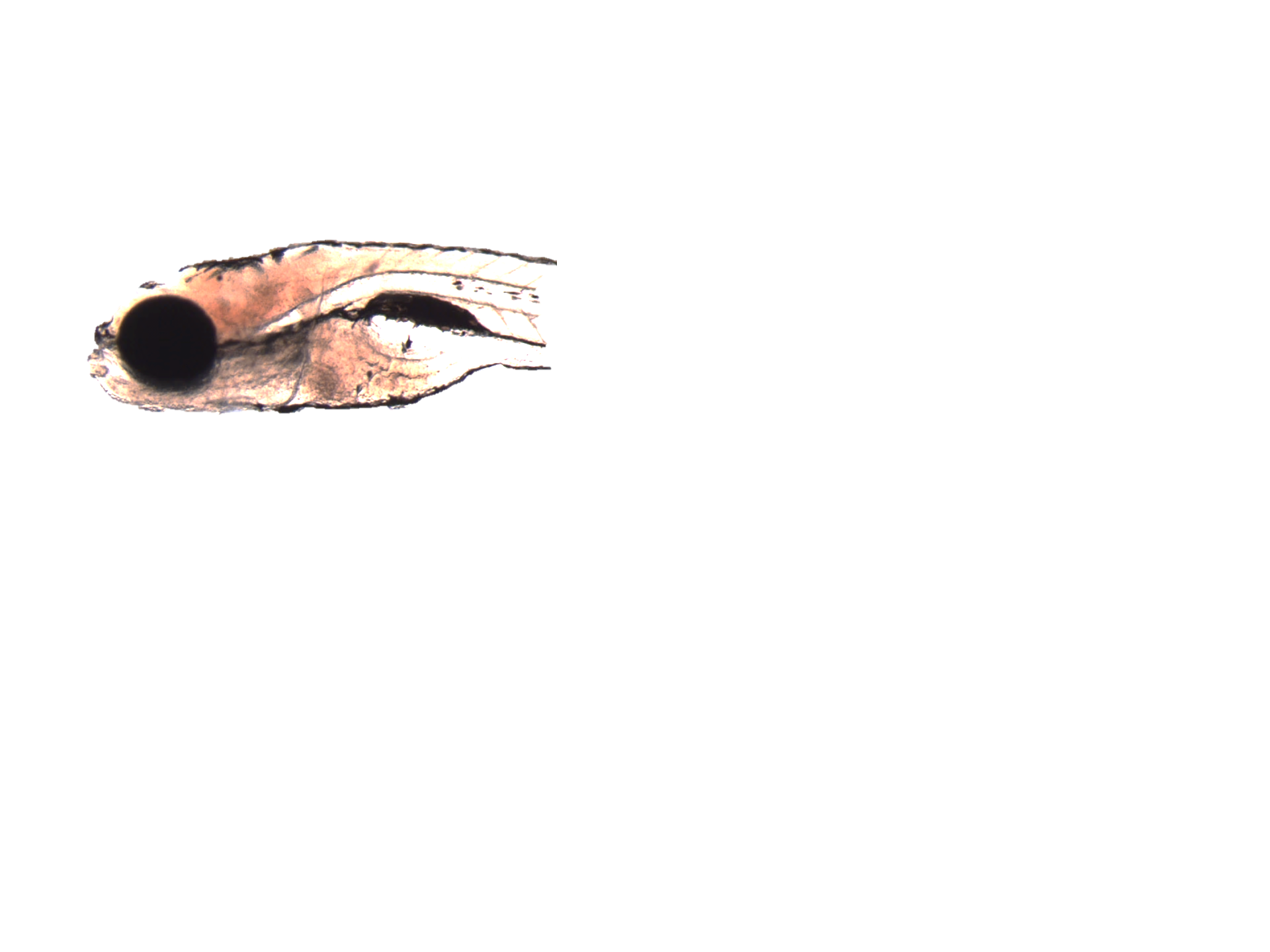

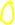

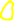

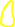

*tm6sf2*

Supplementary Figure 3: Ethanol and HFD increased liver lipid accumulation. ORO staining shows lipid accumulation in pnpla3, faf2 and tm6sf2 crispants challenged with EtOH or high fat diet from. Liver area is marked in yellow line
